# Dosage sensitivity of the loop extrusion rate confers tunability to genome folding while creating vulnerability to genetic disruption

**DOI:** 10.1101/2025.08.14.667581

**Authors:** Rini Shah, Maxime M. C. Tortora, Nessim Louafi, Hadi Rahmaninejad, Karissa L. Hansen, Erika C. Anderson, David Wen, Luca Giorgetti, Geoffrey Fudenberg, Elphège P. Nora

**Affiliations:** Cardiovascular Research Institute, University of California; San Francisco, San Francisco, USA; Department of Quantitative and Computational Biology, University of Southern California, Los Angeles, CA, USA; Friedrich Miescher Institute for Biomedical Research, Basel, Switzerland; Developmental and Stem Cell Biology Graduate Program, University of California, San Francisco, CA, USA; Department of Biochemistry and Biophysics, University of California, San Francisco, San Francisco, CA, USA; Chan-Zuckerberg Biohub San Francisco, CA, USA

## Abstract

Genome folding is not static, but emerges from dynamic processes that control transcription, replication, recombination, and repair. DNA loop extrusion by cohesin is central to genome organization, yet it remains unclear how cells can tune extrusion kinetics to achieve precise and functional chromosome folding patterns. Here we discover extrusion rate acts as a tunable biophysical parameter in cells, quantitatively dialed by the respective dosage of the cohesin cofactors NIPBL and PDS5. Modulation of extrusion rate can offset changes in cohesin lifetime to buffer steady-state chromosome structure and transcriptional states, even in the face of abnormal extrusion dynamics. These findings provide a long-sought mechanistic basis for the genetic interactions between cohesin cofactors and the molecular origin of haploinsufficiency in cohesinopathies, such as Cornelia de Lange syndrome.

## Main text

The physical organization of chromosomes influences countless genomic processes, including transcription, DNA repair, replication, recombination and segregation (*1*). Throughout the domains of life, Structural Maintenance of Chromosome (SMC)-family protein complexes act as key architects of chromosome folding, through their ability to extrude DNA loops (*2–4*). Cohesin has emerged as the main loop extrusion machinery during interphase in vertebrates (*5*, *6*), with several of its cofactors being crucial for proper chromosome morphology (*7–13*) and downstream genomic processes, that range from transcriptional regulation(*14–16*), stochastic choices amongst gene clusters (*17–20*), heterochromatin integrity (*21–23*), as well as DNA recombination, replication and repair (*24–30*).

Importantly, mutations in cohesin often produce distinct phenotypic and clinical outcomes from those in its cofactors (*31*, *32*), suggesting they perturb different molecular functions. For example, we still do not understand why mutations that reduce overall cohesin dosage are rarely observed in Cornelia de Lange syndrome, whereas heterozygous mutations that only reduce NIPBL expression by 30-40% cause profound developmental abnormalities (*33–35*).

While much is known qualitatively about the biochemical roles of cohesin cofactors (fig. 1A), the field still lacks a systems-level understanding of how they act together to translate molecular interactions with cohesin into the large-scale organization and function of the genome. NIPBL, initially identified as a cohesin loader (*36–38*), appears to play broader roles in vivo (*11*, *39*) and is essential for loop extrusion *in vitro* (*40*, *41*); WAPL promotes cohesin unloading (*42*, *43*); PDS5 facilitates WAPL-mediated unloading (*44*) and can also compete with NIPBL for cohesin binding *in vitro* (*45*, *46*). Despite this knowledge, combinatorial perturbations of these regulators often produce phenotypes that are not readily explained by how we understand each cofactor to work in isolation (*10*, *47–49*), limiting our ability to predict or rationalize the contribution of loop extrusion to genome biology *in vivo*. This gap limits our ability to explain why modest alterations in the stoichiometry of cohesin co-factors can cause severe developmental consequences. Addressing this challenge requires mechanistic models that can directly connect quantitative changes in the dynamics of DNA looping to steady-state genome organization and function.

**Fig. 1:**
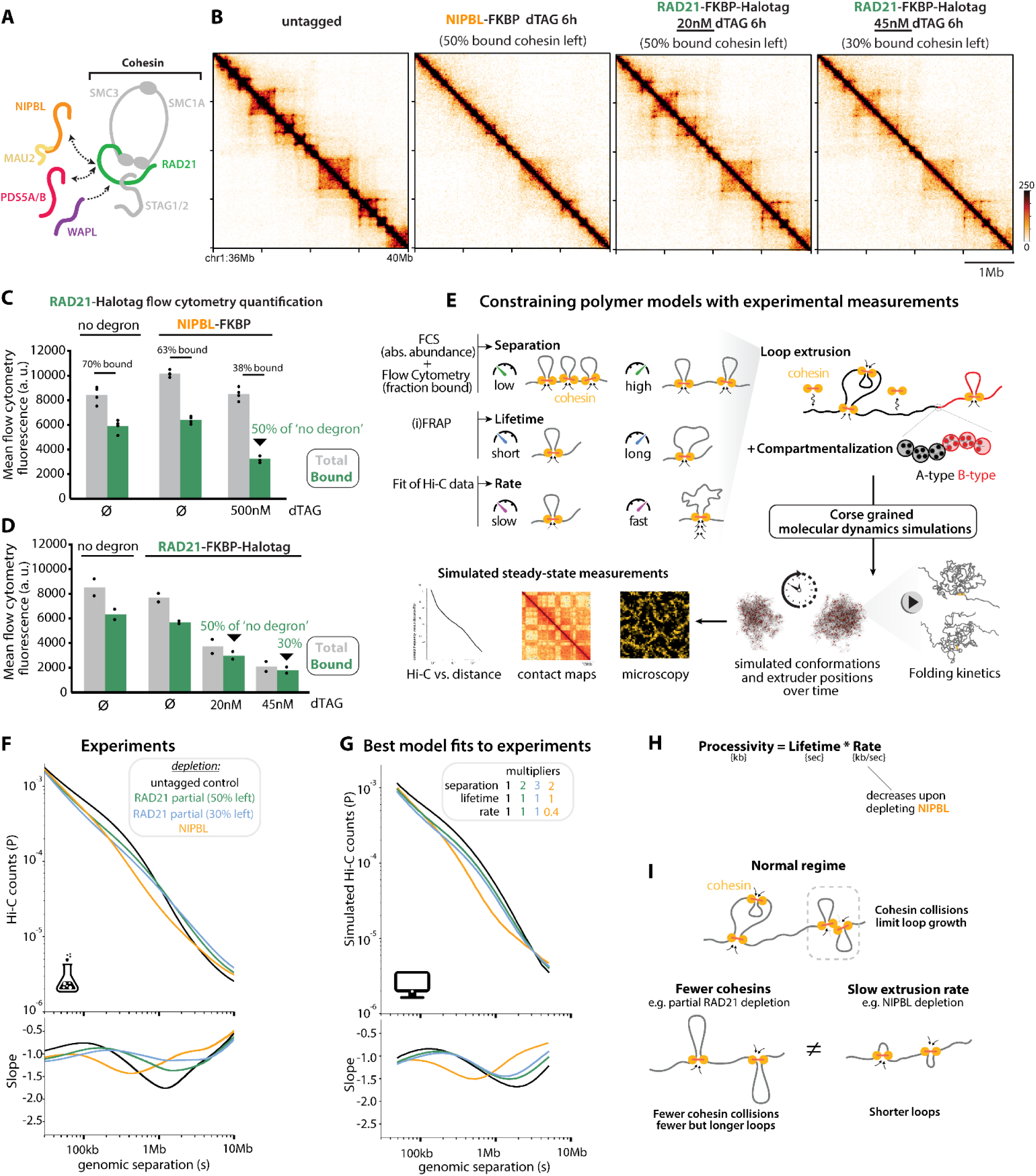
Genome folding defects upon acute NIPBL depletion are best explained by a slower extrusion rate, not by lower cohesin loading. (A) Schematic of cohesin core subunits and cofactors. (B) Hi-C contact maps (20kb bins) after acute NIPBL depletion as in(*14*) or partial cohesin (RAD21) titration with inducible degrons in mouse embryonic stem cells. Reducing cohesin dosage causes distinct Hi-C patterns from NIPBL depletion. (C) Flow cytometry of Halo-tagged RAD21 cells (see Methods). Acute NIPBL depletion only reduces chromatin-bound cohesin 2 fold, even after 1-2 cell divisions (see fig. S2). (D) Flow cytometry quantification of Halo-tagged RAD21 levels after partial induction of the FKBP degron with limiting dTAG amounts, using RAD21-FKBP-Halotag cells. (E) Overview of the polymer modeling approach explaining how extrusion parameters of separation, lifetime, and extrusion rate are calibrated from experimental measurements to reflect the basal state (see fig. S3). (F) Experimental *P(s)* curves from Hi-C after NIPBL or partial RAD21 depletion. (G) Simulated *P(s)* curves corresponding to the parameters that best fit the experiments (see fig. S4 for fitting procedure) (H) Lower cohesin processivity in NIPBL-depleted cells is specifically explained by a reduction in rate. (I) Summary cartoons.

Because properties of loop extrusion are not direct readouts from cellular assays, including Hi-C and microscopy, *in silico* modeling has emerged as a powerful approach to bridge this gap (1, 5). However, such models require fitting multiple parameters that often remain underconstrained by available experimental data. This in turn conflates the inferences for distinct parameters of cohesin extrusion dynamics, obscures the modularity of their control by separate cofactors, and precludes predictions that can be directly tested in cells. Progress in experimental genetic perturbation protocols therefore needs to be coupled with the development of new strategies for modeling and inference. Such integrative approaches are poised to enable unambiguous mapping between cofactor perturbations and individual biophysical parameters of loop extrusion, allowing to gain molecular insight into fundamental genome biology and clinically relevant aspects of cohesin biology.

Here, we systematically determined how individual cohesin cofactors influence distinct biophysical parameters of loop extrusion: the number of loaded cohesins (*separation* between extruders), their residence time on chromatin (*lifetime*), and how fast they extrude (*rate*). We elucidated this molecular control system by combining: (1) an unprecedented suite of degron tags on cohesin and cofactors that enable precisely titrated perturbations in mouse embryonic stem cells (ESCs); (2) calibrated biophysical models capable of using multimodal experimental readouts from Hi-C, microscopy in fixed or live cells, and flow cytometry to infer alterations to parameters of loop extrusion after cellular perturbations. Leveraging this approach, we discovered that the rate of cohesin loop extrusion is a biologically controlled parameter, governed by the cellular balance of the cohesin cofactors NIPBL and PDS5. A tunable loop extrusion rate enables cells to offset changes in other aspects of cohesin dynamics, such as lifetime on DNA. The modularity of this molecular control system enables cells to precisely tune genome organization, from loops to compartments and nuclear architecture, and to finely adjust downstream processes such as transcription. Yet, at the same time, it creates exquisite sensitivity to dosage variation in NIPBL – uncovering a long-sought mechanistic explanation for the haploinsufficiency observed in Cornelia de Lange syndrome.

## Results

### Reduced cohesin loading does not explain chromosome folding defects upon NIPBL depletion

Since NIPBL is needed for cohesin loading, we first investigated whether defects caused by loss of NIPBL can be explained solely by impaired loading. To acutely deplete NIPBL from mouse embryonic stem cells (ESCs), we used a cell line where the endogenous protein can be degraded with the FKBP12^F36V^ degron upon induction by the small molecule dTAG-13 (*14*, *50*) – hereafter abbreviated FKBP and dTAG. In these cells, levels of NIPBL (and its binding partner MAU2) drop below 10% within hours without causing cell cycle defects(*14*). Hi-C defects resembled acute RAD21 degradation with an auxin-inducible degron (AID,fig. 1B and fig. S1) (*14*, *51–54*). These defects arise from impaired cohesin loop extrusion, not sister chromatid cohesion, as contacts between sisters only marginally contribute to bulk Hi-C patterns in mammalian cells - even in G2 (*55*).

To precisely measure the amount of cohesin left on chromatin after NIPBL depletion, we tagged the endogenous core cohesin subunit RAD21 with a Halotag (which had no effect on chromosome folding, fig. S1B) in the NIPBL-FKBP cells. We then pre-extracted soluble proteins from the cells and precisely quantified the remaining chromatin-bound cohesin by flow-cytometry (fig. S2A, Methods). With this highly quantitative approach (56), we estimated that ∼60-70% of RAD21 molecules are bound to chromatin in WT ESCs (fig. 1C), in line with previous orthogonal estimates from live imaging and Western blot across a variety of mammalian cell types (*39*, *56–59*). NIPBL depletion only reduced the abundance of chromatin-bound cohesin two-fold (fig. 1C) even after 24h (two full cell cycles, figure S2B), consistent with previous RNAi experiments (*7*).

To assess whether the two-fold reduction in chromatin-bound cohesin accounts for the chromosome folding defects observed upon NIPBL depletion, we engineered cells in which cohesin abundance can be directly reduced. We created RAD21-FKBP-Halotag cells and titrated dTAG concentrations to achieve precise intermediate levels of bound RAD21 that we quantified by flow cytometry (fig. 1D and fig. S2C). The dramatic Hi-C defects caused by NIPBL depletion were not recapitulated by reducing chromatin-bound cohesin levels two- or even three-fold (fig. 1B, conditions that do not induce confounding cell cycle defects, fig. S1E).

Acute NIPBL depletion therefore disrupts chromosome folding more severely than expected from the remaining amount of chromatin-bound cohesin. Thus, NIPBL must control chromosome folding through functions beyond simply loading cohesin onto chromatin.

### Calibrated biophysical simulations connect extrusion kinetics to genome folding

If not through reduced cohesin loading, NIPBL depletion could in principle attenuate long-range chromatin interactions by lowering extrusion processivity, the distance traveled by an unimpeded cohesin. Processivity depends both on lifetime, the time cohesin spends on chromatin during extrusion, and on extrusion rate, the average speed at which cohesin extrudes. Inverse fluorescence recovery after photobleaching (iFRAP) indicated that cohesin residence time does not measurably change upon NIPBL depletion (fig. S2E). This leaves the hypothesis that, beyond loading, NIPBL abundance may modulate the rate of loop extrusion.

To gain quantitative insight into how lowering the extrusion rate would affect genome folding, we developed a modeling approach that explicitly enables modulating rate independently of lifetime (fig. 1E, Methods). This is distinct from earlier implementations that modulated the relaxation rate of the polymer fiber but did not consider how increased stepping rates of extruders would translate into larger distances traveled along the chromatin fiber in a given amount of time (*60*, *61*). Importantly, this enables us to study the respective contributions of lifetime and extrusion rate to processivity. Similar to previous implementations (*60*, *62–64*), we coupled a 1D lattice model of loop extruder dynamics with a 3D polymer model to simulate 50 Mb of chromatin (fig. S3A). We then parameterized the basal extrusion state based on available mESC data. We modeled A/B compartmentalization as a block copolymer that mirrors the arrangement of active and inactive regions along an endogenous chromosome. We set the separation between extruders based on absolute cohesin abundance previously measured by Fluorescence Correlation Spectroscopy (FCS) (*65*) (fig. S3B). We set the extruder lifetime based on FRAP data (*56*), and the correspondence between simulated and experimental time based on locus tracking data (*66*, *67*). To estimate the rate of extrusion, for which direct *in vivo* measurements are currently lacking, we fit simulations to the experimental Hi-C *P(s)* from untagged wild-type cells (fig. S3E-F and Methods). The resulting model for WT untagged cells corresponds to an average separation of 185 kb between extruders, a lifetime of 22 minutes, and average extrusion rate of 0.25 kb/sec.

### Polymer simulations support a prominent role for NIPBL beyond cohesin loading in controlling loop extrusion rate

Using simulations, we investigated what changes in extrusion parameters could account for the chromosome folding defects we observed after NIPBL depletion. Experimental *P(s)* curves from NIPBL-depleted (with NIPBL-FKBP) or partially RAD21-depleted cells (with RAD21-FKBP-Halotag) both displayed reduced contact frequencies between ∼20 kb-2 Mb (fig. 1F). With polymer simulations, we then assessed how *P(s)* curves change with decreasing numbers of extruders, equivalent to increasing the genomic separation between extruders (fig. S4A). Clear, though subtle, differences were observed in simulations across the range of extruder separations considered. We thus considered the goodness-of-fit between simulated and experimental *P(s)* curves to determine the best-fitting simulation parameters. Remarkably, the best fit to the experimental *P(s)* profile from cells with chromatin-bound RAD21 reduced to 50% was achieved in simulations with 2-fold increased extruder separation (fig. S4B). Similarly, the best fit to the experimental *P(s)* profiles from cells with chromatin-bound RAD21 reduced to 30% was achieved with simulations with 3-fold increased extruder separation. Therefore, this fitting approach between experimental and simulated data can account for subtle changes in *P(s)* by correctly incriminating the loop extrusion parameter corresponding to the number of extruders (compare experiments to best fitting simulations in fig. 2F-G).

**Fig. 2:**
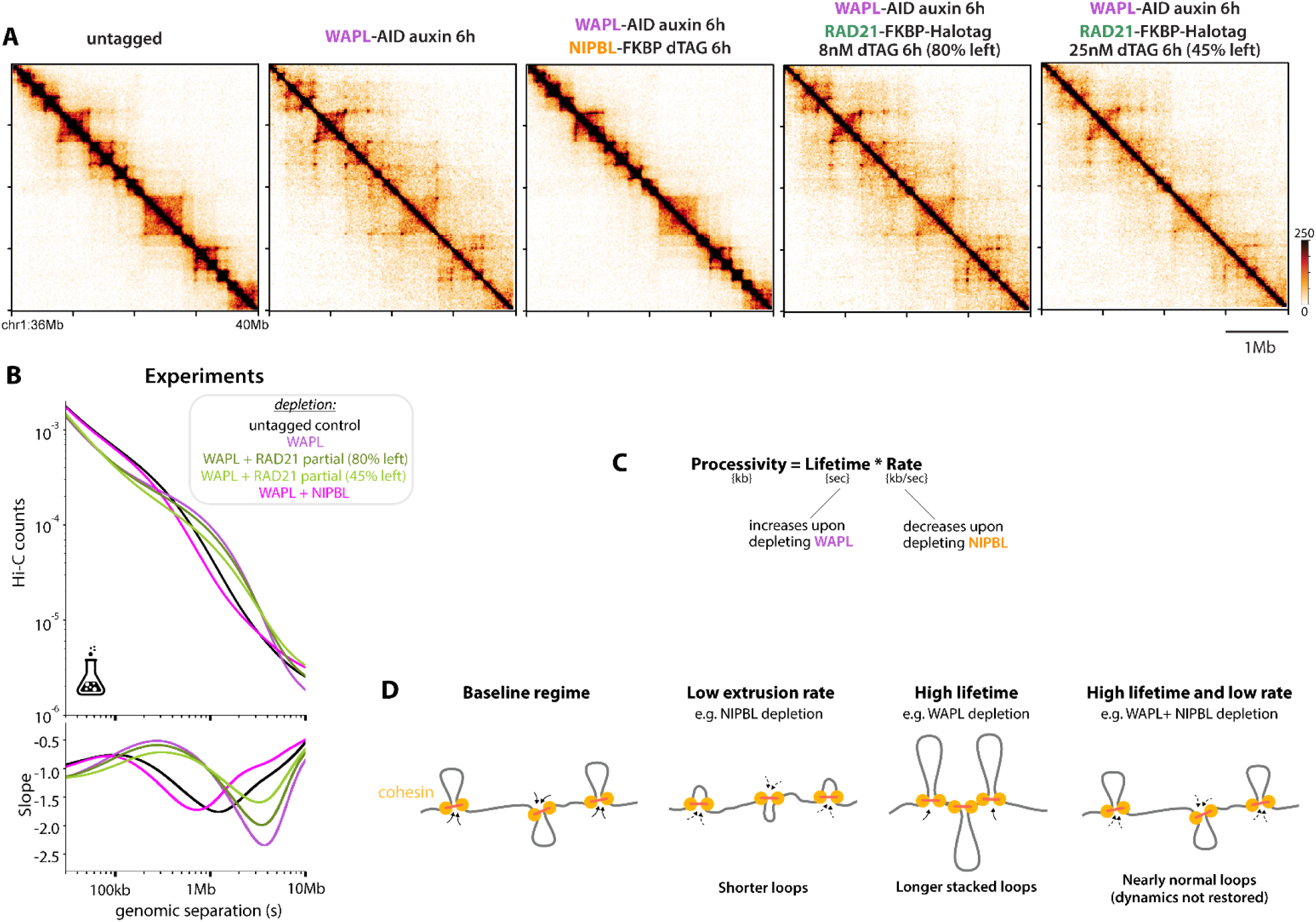
Reduced extrusion rate after NIPBL co-depletion, not reduced loading, mitigates genome misfolding caused by increased lifetime upon WAPL inactivation. (A and B) Experimental Hi-C (A) and corresponding *P(s)* curves showing that chromosome folding after acute co-depletion of WAPL + NIPBL resembles that of normal cells, but is distinct from patterns observed with WAPL + partial RAD21 co-depletion at equivalent amount of bound cohesin (see fig. S6). (C) Considering that processivity is the product of lifetime and rate provides an immediate explanation for how NIPBL depletion (which reduces extrusion rate, see Fig. 1) can counteract the effect of WAPL depletion (which increases lifetime). (B) (D) Summary cartoons.

We next explored the effect of decreasing extrusion rates on *P(s)* in simulations. We observed that lowering extrusion rates had much greater effects than increasing cohesin separation (fig. S4A). Although NIPBL-depleted cells have 2-fold less bound cohesin (fig. 1C), increasing extruder separation alone yielded poor fits to the experimental Hi-C data (fig. S4B). In contrast, lowering extrusion rates in the context of 50% chromatin-bound cohesin provided an excellent fit to the experimental NIPBL Hi-C, with best agreement obtained at 40-50% of initial extrusion rates (fig. S4C). Further supporting our inference, varying extrusion rate alone offered poor fits to the partially depleted RAD21 datasets (fig. S4C). Simulations thus provide quantitative evidence that the Hi-C defects after NIPBL depletion cannot be explained by reduced cohesin loading alone, and instead suggest that NIPBL abundance could shape chromosome folding by modulating the rate of loop extrusion, thereby accounting for its contribution to cohesin processivity (fig. 1H).

Qualitative support for the differential impact of NIPBL versus RAD21 depletion comes from considering how the inflection point in *P(s)* shifts in these two conditions (fig. S5A). This feature of *P(s)*, termed s[P’max], relates to the average length of loops when they are sufficiently separated (*68*). For NIPBL depletion, s[P’max] decreased (from ∼90 kb to ∼55 kb). In contrast, after partial cohesin depletion, s[P’max] increased (up to ∼150 kb for 50% remaining bound cohesin, or ∼210 kb for 30%, fig. S5A). These distinct effects of NIPBL vs. partial cohesin depletions on s[P’max] were reproduced in simulations (fig. 1G). By monitoring the location of extruders on simulated polymers we observed that, in the baseline regime, nearly 70% of extruders have collided on at least one side (fig. S5B). At higher extruder separations, collisions are reduced and average loop lengths increased (fig. S5C), explaining how depleting RAD21 led to a rightward shift in the *P(s)* inflection point (fig. 1F-G).

Altogether, simulations and experiments indicate that cohesin collisions limit loop growth in the normal regime (fig. 1H). Lowering cohesin abundance with partial RAD21 depletion increases the separation between extruders and enables loops to grow slightly larger. Therefore, the *P(s)* measured after NIPBL depletion cannot be explained by reduced cohesin loading. These observations further support the idea that, in addition to the reduction of bound cohesin levels, NIPBL-depleted cells experience a lower rate of extrusion.

### Reduced extrusion rate after NIPBL depletion, not reduced cohesin loading, can counteract the increase in cohesin lifetime caused by WAPL inactivation

A role for NIPBL in setting the rate of loop extrusion yields testable predictions about how it interacts genetically with cofactors controlling other steps of the extrusion process. Several studies have reported that impairing NIPBL functions can prevent the 3D genome defects observed upon WAPL inactivation (*10*, *47*, *69*). WAPL unloads cohesin and thereby controls lifetime (*37*, *42*, *43*). Yet, NIPBL co-depletion does not restore cohesin lifetime (*10*). How, then, do NIPBL and WAPL depletions offset each other in chromosome folding? To distinguish whether compensation arises from reduced extrusion rate or decreased cohesin loading, we engineered WAPL-AID ESCs (*53*, *70*) to either co-deplete NIPBL or partially co-deplete RAD21. As previously reported in other cell types, Hi-C patterns and *P(s)* curves from cells acutely co-depleted for WAPL and NIPBL closely mirrored those of undepleted cells (fig. 2A and 2B). In contrast, simultaneously depleting WAPL and reducing RAD21 levels did not cause Hi-C patterns to resemble those of undepleted cells, even when titrated to precisely match the levels obtained after NIPBL co-depletion (fig. 2A, 2B and fig. S6). This indicates that NIPBL co-depletion mitigates WAPL inactivation through means other than cohesin loading.

Viewing NIPBL as controlling the rate of loop extrusion immediately clarifies how its co-depletion counteracts the genome folding defects caused by WAPL inactivation, in spite of cohesin lifetime remaining aberrantly high. Indeed, given that processivity is the product of lifetime and rate (fig. 2C), the decrease in loop extrusion rate from NIPBL depletion offsets the increase in lifetime from WAPL depletion (fig. 2D), thereby preserving the size of extruded loops.

Beyond ascertaining the role of NIPBL in controlling the rate of loop extrusion, these results provide a long-sought mechanistic explanation for the genetic interaction between NIPBL and WAPL. Therefore, opposing changes to distinct extrusion parameters can counterbalance one another to buffer chromosome architecture in the face of shifts in cohesin dynamics, delineating the existence and principles of a flexible control mechanism for genome organization.

### Modeling of Hi-C indicates that PDS5 depletion accelerates loop extrusion rates

We next considered the role of cellular PDS5 in loop extrusion. PDS5 is a protein that binds cohesin and promotes WAPL-mediated cohesin unloading (*71–73*). Yet PDS5-depleted cells do not exhibit Hi-C patterns identical to WAPL-depleted cells, indicating that PDS5 has functions in genome folding beyond mediating unloading (*7*, *48*, *74*, *75*). To gain further insight, we performed Hi-C in cells with near-complete depletion of both PDS5A and PDS5B (hereafter PDS5AB) for 6h using the AID system, as we observed that both paralogs contribute to genome folding in ESCs (fig. S7A-D). In line with previous reports in other cell types (*7*, *74*), PDS5AB depletion resulted in more altered *P(s)*, and less sharply defined Hi-C dots compared to WAPL depletion (fig. 3A-C). Consistent with unloading defects, both WAPL and PDS5AB depletion increase the amount of chromatin-bound cohesin (fig. S7E). These results are not solely driven by cell cycle defects, as short-term depletion of WAPL in ESCs does not affect the cell cycle (*53*), nor does PDS5AB depletion (fig. S7F). Longer PDS5AB depletion caused S-phase stress, as previously reported (*25*, *76*), but only beyond 24h.

**Fig. 3:**
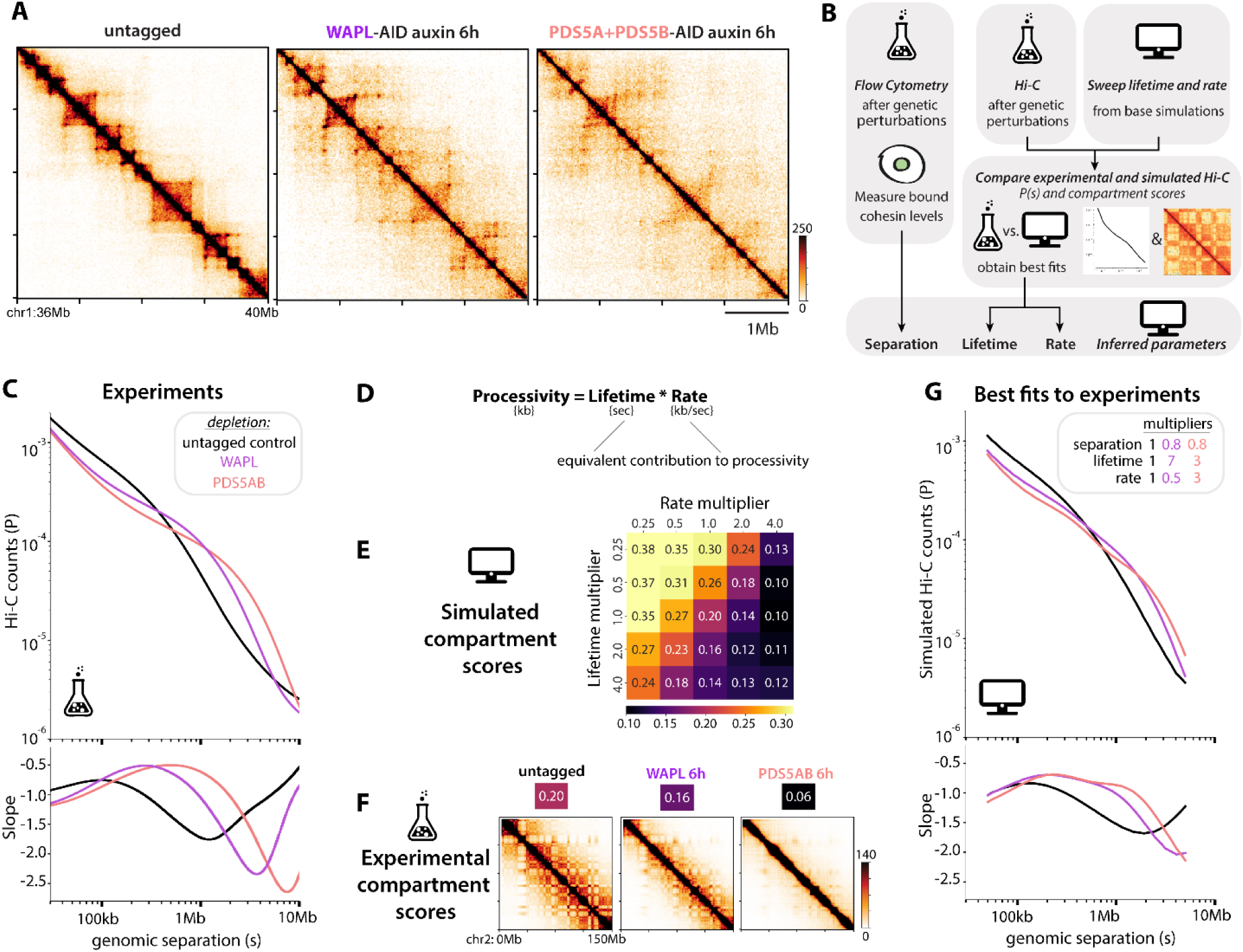
Compartment scores disambiguate the contribution of cohesin lifetime and extrusion rate to genome misfolding upon WAPL or PDS5AB depletion. (A) Hi-C after acute depletion of WAPL or combined PDS5A and PDS5B (hereafter abbreviated PDS5AB). (B) Workflow of the experimentally calibrated biophysical framework to measure separation and infer lifetime and rate parameters. (C) Experimental *P(s)* curves highlighting the greater impact of depleting PDS5AB than WAPL. (D) The equivalent contribution of lifetime and rate to processivity creates ambiguity when trying to infer their respective contribution to *P(s)*. (E) Simulations reveal that compartment scores decrease more steeply upon increasing extrusion rate than increasing cohesin lifetime. Lower scores indicate weaker separation of A and B compartments. Here base extruder separation was used for all simulations. (F) Experimental Hi-C contact maps (chromosome-wide, 1 Mb bins) and corresponding compartment scores (for the same chr2:66-116Mb region used in simulations) are more altered upon depleting PDS5AB than WAPL. (G) Simulated *P(s)* with multipliers chosen from fitting both *P(s)* and compartment scores (note 0.8x cohesin separation comes from calibration to the experimentally-measured amount of chromatin-bound RAD21, see fig. S7-8).

To elucidate how WAPL and PDS5AB depletion differentially impact genome folding, we approached this challenge as a system of three unknowns - loaded cohesin number, extrusion rate, and lifetime - that we solved by quantitatively estimating three metrics experimentally after each perturbation: flow cytometry for chromatin-bound cohesin, and *P(s)* and compartment scores from Hi-C (fig. 3B). Since we calibrated the number of extruders in the simulations to the experimentally measured amount of chromatin-bound cohesin, we could focus on estimating lifetime and extrusion rate from Hi-C. Importantly, as *P(s)* is largely influenced by the extruder processivity (*i.e*. the product of lifetime and rate), this metric alone is unable to distinguish a contribution of higher lifetime from higher extrusion rate (fig. 3D). Indeed, when we increased cohesin lifetime and extrusion rate either in isolation or in combination, and compared each simulated *P(s)* to the experimental Hi-C after WAPL or PDS5AB depletion, we obtained a large number of combinations with excellent fits (fig. S8). We therefore turned to additional metrics beyond *P(s)* to disambiguate the contribution of lifetime and rate to chromosome folding patterns.

We noticed that, in simulations, increasing extrusion rate reduced the segregation of A and B compartments more effectively than increasing extruder lifetimes (fig. 3E). Compartmentalization scores therefore enabled us to disambiguate the contribution of cohesin lifetime versus extrusion rate to processivity, and quantitatively estimate the extent to which these were differentially after PDS5AB versus WAPL depletion. Experimentally, PDS5AB depletion lowered compartment scores further than WAPL depletion (fig. 3F). Requiring a joint best-fit to both experimental *P(s)* and compartment scores after WAPL and PDSAB depletion led to a unique set of best fitting parameters for each condition. PDS5AB depletion only increases cohesin lifetime three-fold (about half of WAPL depletion), but also increases extrusion rates three-fold (fig. 3G). These observations indicate that, even though cohesin lifetime is less perturbed by lowering PDS5AB dosage than WAPL dosage, the increase in extrusion rate drives more profound changes in genome folding.

### Microscopy-based evidence that PDS5 depletion does not fully disable unloading but accelerates loop extrusion

We next tested predictions of the PDS5AB and WAPL simulations by considering orthogonal manifestations of loop extrusion that were not used to inform the selection of model parameters. Disabling unloading causes extruding cohesins to collide and form axial structures known as vermicelli (*8*, *77*), visible as spatial clusters of cohesins in the nucleus. We therefore analyzed the location of extruders in individual simulated conformations, performing *in silico* microscopy, using the parameters for lifetime and extrusion rate that we previously identified as best fitting the Hi-C data. While simulations of WAPL depletion exhibited some clustering of extruders, simulations of PDS5AB depletion displayed more pronounced axial structures (fig. 4A). We also observed this pattern experimentally using fixed-cell microscopy: PDS5AB depleted cells displayed stronger cohesin clustering than WAPL depleted cells (fig. 4B). Microscopy thus argues that cells depleted for PDS5AB experience faster extrusion rates than WAPL depleted cells, supporting the distinct roles of these cofactors we deduced from fitting simulations to Hi-C.

**Fig. 4:**
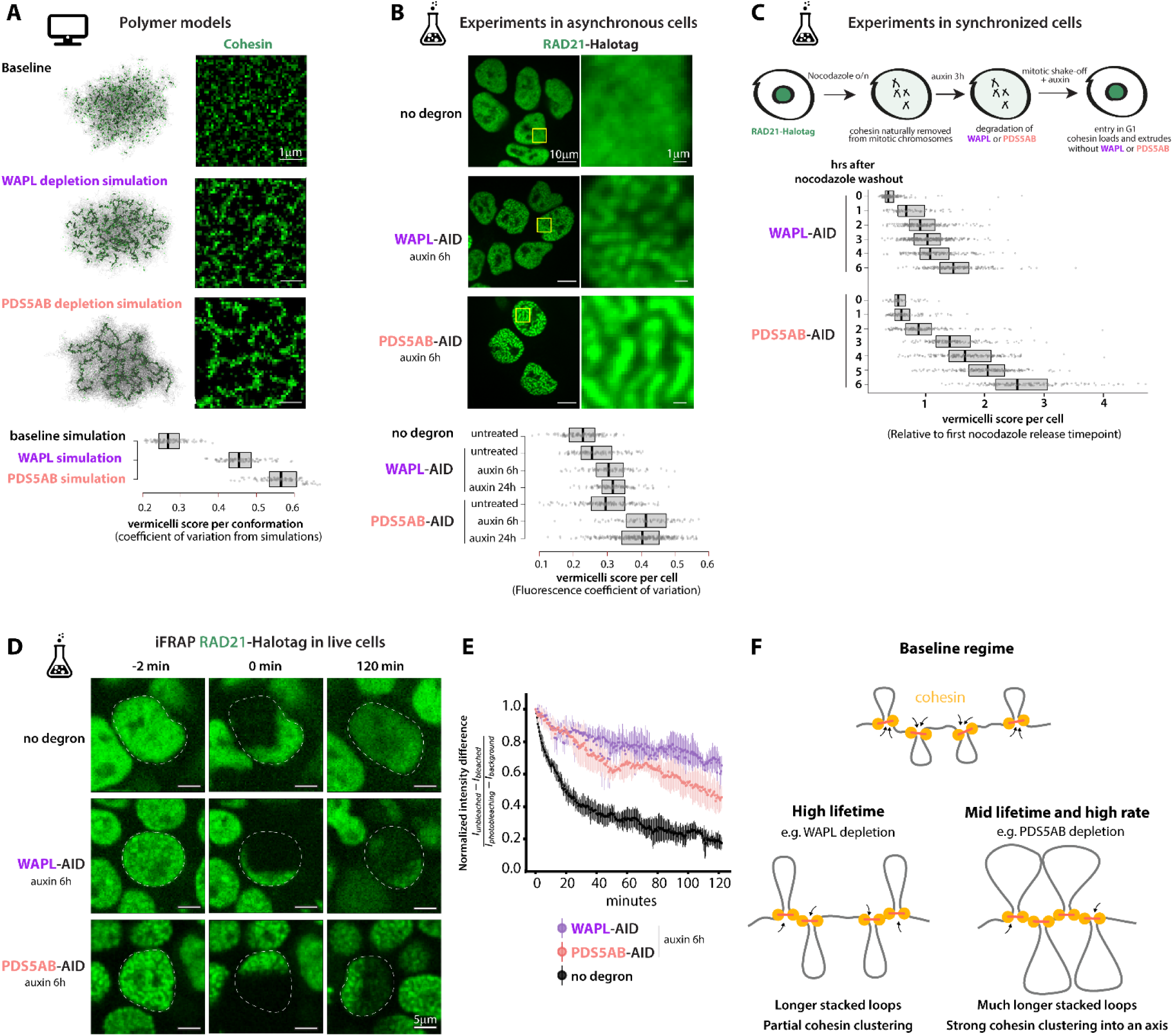
PDS5AB depletion accelerates the rate of cohesin loop extrusion. (A) Example of simulated chromosomes (left) and *in silico* microscopy images obtained from analyzing the location of extruders (right). Simulations were conducted using extrusion parameters that best fit Hi-C experiments (see fig. 3). The models predict that cohesin clusters more strongly into axial structures (vermicelli) upon PDS5AB depletion compared to WAPL depletion. (B) Experimental staining of RAD21 in Halo-tagged subclones (JFX646 dye) highlighting stronger clustering into vermicelli after PDS5AB depletion than WAPL depletion. Vermicelli scores correspond to the coefficient of variation in (simulated) microscopy images (see Methods). (C) Cohesin clustering into vermicelli occurs more rapidly after G1 entry following PDS5AB depletion than WAPL depletion, consistent with PDS5AB depletion accelerating the rate of loop extrusion. (D) Inverse Fluorescence Recovery After Photobleaching (iFRAP) of RAD21-Halotag after either WAPL or PDS5AB depletion. (E) iFRAP quantifications showing that depleting PDS5AB does not increase cohesin lifetime as much as depleting WAPL. n=12-18 cells, in support of the model predictions. Shown are medians +/- standard deviation (see fig. S9). (F) Summary cartoons.

To more directly ascertain that PDS5AB depletion increases the rate of cohesin extrusion, we assayed the speed of vermicelli formation in cells. Cohesin is naturally removed from chromosomes in mitosis through the prophase pathway, only initiating extrusion at the onset of G1 (*78*). If chromosome extrusion is faster after PDS5AB than WAPL depletion, then compaction into vermicelli structures in early G1 should also occur faster after PDS5AB depletion. To test this, we used nocodazole to synchronize ESCs in M phase (fig. 4C, S9A-B) and depleted either WAPL or PDS5AB for three hours by adding auxin. We then collected mitotic cells by mitotic shake-off, and released cells into G1 by washing off nocodazole (keeping auxin in the medium to suppress either WAPL or PDS5AB expression). A microscopy time-course over six hours after release into G1 revealed that vermicelli form faster and end up more pronounced in the absence of PDS5AB than in the absence of WAPL (fig. 4C). These experiments therefore support that PDS5AB depletion accelerates the rate of loop extrusion, and rule out possible technical confounders in asynchronous cells, such as eventual differences in cell cycle length or degron leakiness between cell lines.

We then tested the model prediction that WAPL depletion leads to a greater increase in cohesin lifetime than PDS5AB depletion using iFRAP for RAD21-Halotag in live ESCs. We observed that PDS5AB depletion increased the residence time of cohesin, but not as much as WAPL depletion (fig. 4D-E, S9C-D), in line with RNAi experiments in HeLa cells (*7*). iFRAP thus quantitatively supports the parameters inferred from our calibrated polymer simulations and Hi-C fitting: while cohesin lifetime increases upon PDS5AB depletion, it does not reach the levels seen after WAPL depletion.

Altogether, these observations ascertain that PDS5 limits both rate of loop extrusion and the lifetime of loop extruders, accounting for its unique role in genome folding, distinct from WAPL. The combination of higher extrusion rates and lifetime explain the Hi-C *P(s)* observed upon PDS5AB depletion, as well as the stronger nuclear clustering of cohesin seen by microscopy (fig. 4F). These results also demonstrate how polymer simulations, although only instructed by Hi-C data and cohesin abundance from flow-cytometry data, can both quantitatively predict meso-scale chromosome organization observable by microscopy, and assign specific molecular roles in extrusion kinetics to cohesin cofactors.

### The NIPBL:PDS5 balance shapes chromosome folding by tuning the rate of cohesin extrusion

We next explored whether the interplay of multiple cohesin cofactors renders rate a tunable property of the extrusion system. We reasoned that the opposing effect of depleting PDS5AB or NIPBL on extrusion rate should offset one another when combined. However, because PDS5AB depletion also increases cohesin lifetime, PDS5AB+NIPBL co-depleted cells should still be partly deficient in unloading. Therefore, the expectation is that Hi-C in PDS5AB+NIPBL co-depleted cells would resemble WAPL-depleted cells, which have increased lifetime. This is indeed what we observed experimentally (fig. 5A-B). This offset is not due to how NIPBL depletion reduces cohesin loading: in the background of PDSA5AB depletion, titrating down RAD21 levels did not create Hi-C patterns reminiscent of WAPL inactivation (fig. 5A-B and fig. S10A). Instead, it shifted the *P(s)* inflection point to substantially higher genomic distances, consistent with a decrease in cohesin-cohesin collisions. As such collisions normally constrain loop growth (fig. 5), fewer collisions would enable loops to extend further upon PDS5AB depletion.

**Fig. 5:**
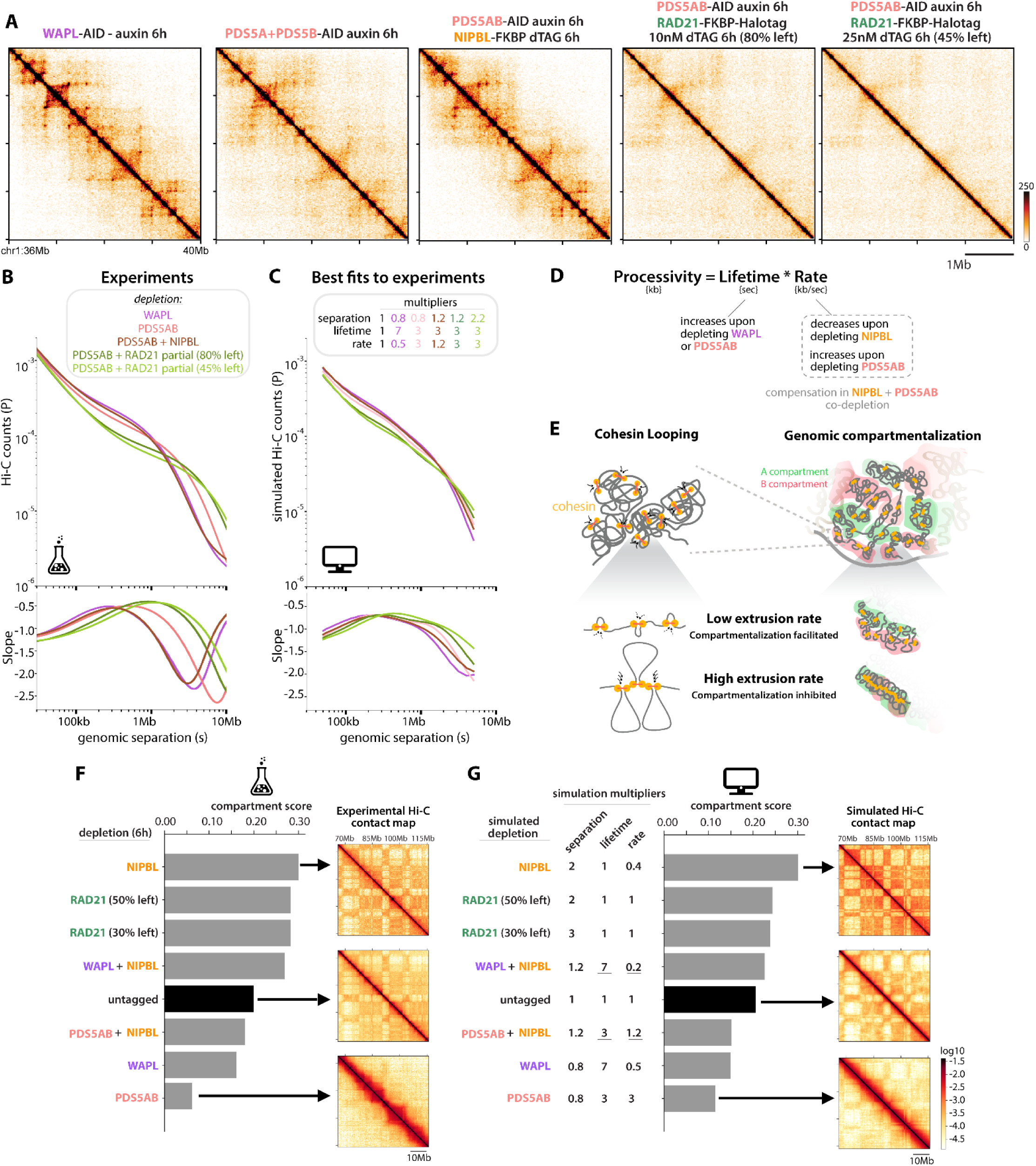
The NIPBL:PDS5 balance shapes chromosome folding across scales by setting the rate of cohesin extrusion. (A and B) Experimental Hi-C (A) and corresponding *P(s)* curves (B) after acute co-depletion of PDS5AB + NIPBL closely resemble WAPL depletion, and are distinct from PDS5AB + partial RAD21 co-depletion. (C) Simulated *P(s)* curves and corresponding parameters of loop extrusion. Rate multipliers for the single NIPBL depletion were obtained as the best fit to experimental *P(s)*, in fig. 1. Lifetime and rate multipliers for PDS5AB-only depletion were obtained as the joint best fit to experimental *P(s)* and compartment scores, in fig. 3. Multipliers for co-depletions were computed by multiplying loop extrusion parameters of the single mutants. Separation multipliers in simulations were calibrated to match experimental measurements of chromatin-bound RAD21 by flow cytometry in each of the single and co-depletions (fig. S10). Simulations indicate that reducing rate (NIPBL depletion) can offset the increase in rate caused by PDS5AB depletion, but not the concomitant increase in lifetime - explaining why NIPBL+PDS5AB *P(s)* closely match that of WAPL (D) Depleting PDS5AB increases processivity by increasing both cohesin lifetime and extrusion rates. Lowering extrusion rate by co-depleting NIPBL and PDS5AB offsets the extrusion rate but does not restore lifetime. Considering the separate effect of PDS5AB depletion on cohesin lifetime and extrusion rate therefore explains why co-depleting NIPBL + PDS5AB leads to similar genome folding patterns as depleting WAPL alone. (E) Illustration of how extrusion rate alters genomic compartmentalization. (F) Ordering of experimental depletions by compartment scores (see fig. S11). (G) Ordering of simulated depletions by compartment scores. Separation values correspond to the experimentally measured chromatin-bound RAD21; the lifetime and rate multipliers for co-depletions (underlined) are the product of the best fits for single depletions, as in fig. 1, 2 and 3. Note the identical ordering of compartment scores between simulations and experiments.

To understand the complex chromosome folding defects that result from disrupting two cohesin cofactors, we simulated each co-depletion. We assigned cohesin separation multipliers using flow cytometry measurements of chromatin-bound RAD21 (Fig. S10A). We set lifetime and extrusion rate by multiplying the parameters from each of the two single depletions (from Figures 2 and 3). The resulting simulated *P(s)* for co-depletions mirrored those for experimental data, supporting the mechanistic explanations we provide for individual co-factors: in PDS5AB+NIPBL co-depletion, the effect of NIPBL depletion lowering extrusion rate is counteracted by PDS5AB depletion raising extrusion rate and lifetime. As a result, after PDS5AB+NIPBL co-depletion rate is re-balanced but lifetime remains elevated, explaining the close resemblance to WAPL depletion (fig. 5B-C). In contrast, reducing cohesin levels by partial depletion of RAD21 cannot compensate for the increase of rate and lifetime caused by PDS5AB depletion.

We reached similar conclusions by modeling co-depletions with WAPL: simulations used by combining the parameters inferred from single depletions (fig. S10B) also matched experimental co-depletion data (fig. 2B). Increasing lifetime (due to WAPL depletion) while reducing extrusion rate (due to NIPBL depletion) results in normal processivity and therefore a *P(s)* curve similar to WT untagged cells. In contrast, RAD21 partial depletion does not compensate for the longer lifetime in WAPL depletion (fig. S10B).

### Rate modulation provides a way to shape genomic compartmentalization, and can buffer the effect of lifetime changes independently of cohesin abundance

To evaluate the generalizability of our models, we examined their predictions for compartmentalization (fig. 5E). All simulations used experimentally-determined cohesin separation, obtained as above by flow cytometry of chromatin-bound RAD21. Remarkably, the ranking of compartment scores across experimental depletions (fig. 5F) was identical to the ranking of simulations (fig. 5G and fig. S11A-C, aside from partial co-depletions of RAD21 on WAPL and PDS5AB backgrounds). Indeed, PDS5AB+NIPBL codepletion had higher compartmentalization than WAPL alone, and WAPL+NIPBL co-depletion had higher compartmentalization than even wild type cells, both in simulations and experiments.

Importantly, reducing the extrusion rate in PDS5AB mutants, via NIPBL co-depletion, greatly restored compartment strength across genomic distances in *cis* and in *trans* (fig. S11D). While partial RAD21 depletion increased compartmentalization in simulations on the background of PDS5AB or WAPL depletion, experimental compartmentalization was enhanced more than predicted for both of these scenarios. We speculate that this stems from the greater relative availability of these residual subunits when cohesin is simultaneously downregulated. Indeed, the higher PDS5AB:RAD21 and WAPL:RAD21 ratios in these co-depleted cell lines could help rescue cohesin lifetimes and extrusion rates towards their basal values. This effect is neglected by the current assumption that the net changes in extrusion parameters induced by simultaneous co-depletions are strictly multiplicative (fig. S11C).

Altogether, our findings elucidate how the modulation of extrusion rate by the NIPBL:PDS5 finely tunes genome folding pattern, from DNA looping to genomic compartmentalization and nuclear organization (fig. 5E), and showcases the broad applicability of our approach to infer molecular dynamics from snapshot data.

### The rate of cohesin extrusion impacts transcriptional states and scales with NIPBL dosage

To explore the relevance of rate modulation to downstream genomic processes, we tested whether Hi-C defects tracked with overall transcriptional changes by performing RNA-seq across six depletion conditions, including our previously reported NIPBL degradation (*14*). We profiled transcriptomes after 1 day of depletion to capture enough dysregulated genes to compare cellular states between perturbations, refraining from correlating transcriptional dysregulations with local genomic features, as the secondary defects would confound such analyses. Samples with similar contacts versus distance curves (as reflected by their relative *P(s)* inflection points, or Hi-C s[P’max] values, see fig. S5A) displayed similar transcriptomes (as estimated by PC1 from principal component analysis, fig. 6A and 6B). Notably, NIPBL+WAPL co-depletion not only produced Hi-C patterns resembling those of untagged cells, but also led to clustering of RNA-seq profiles with untagged controls. Similarly, NIPBL+PDS5AB depletion and WAPL depletion, which caused similar Hi-C patterns, led to similar transcriptomic profiles. PDS5AB depletion, which caused the most significant change in processivity, also induced the greatest overall transcriptomic alterations. These findings illustrate that changes in loop extrusion kinetics not only affect 3D genome folding but can also have a profound impact on overall transcriptional states, providing molecular insight into how genetic interactions between cohesin cofactors can correct transcriptomes disrupted by alterations in specific loop extrusion parameters.

**Fig. 6:**
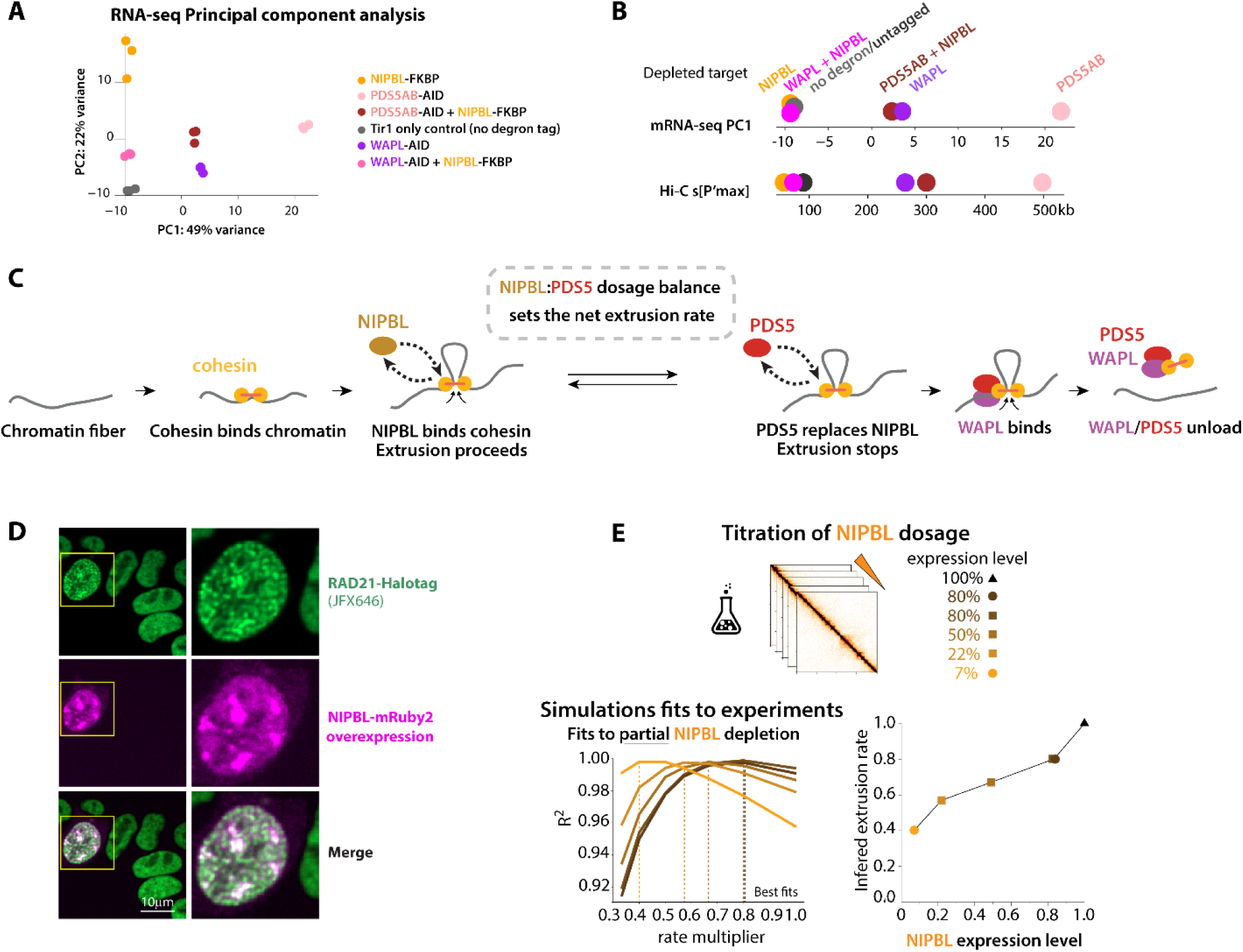
The rate of loop extrusion affects transcriptional states and is quantitatively tuned by the dosage of NIPBL in cells. (A) RNA-seq PCA in the indicated genotypes treated with auxin or dTAG or 1d, showing PC1 and PC2 and all three replicates separately. Tir1 are control cells expressing the auxin receptor transgene but without any degron knockin. Transcriptomes of NIPBL-FKBP cells were obtained from Hansen et al.(*14*). (B) *top:* RNA-seq Principal component analysis PC1 values separate depletion conditions according to their overall transcriptomic similarity, averaging replicates. *Bottom:* Samples ordered by genomic distance of their inflection points derived from Hi-C *P(s)* curves (s[P’max], which relates to loop size - fig. S5). Samples with similar Hi-C s[P’max] have similar transcriptomes. (C) Cartoon model illustrating how the NIPBL:PDS5 balance sets the net rate of cohesin loop extrusion. (D) Transient NIPBL over-expression in RAD21-Halotag ESCs is sufficient to cause chromosome compaction into vermicelli. Inset shows a representative cell with high NIPBL expression. (E) Approach to determine how extrusion rates scale with NIPBL dosage: (top) Hi-C maps were collected at four levels of experimentally-titrated NIPBL dosages; (bottom left) best fits to simulations were computed to infer relative extrusion rates (as a multiplier relative to baseline untagged cells - 0.25kb/s). (bottom right) inferred extrusion rates were plotted as a function of NIPBL expression levels (circles: data points obtained with NIPBL-FKBP cells; squares: NIPBL-FKBP-GFP cells; triangle: untagged control cells). Bound cohesin levels were measured with RAD21-Halotag subclones (see fig. S12).

Taken together, our conclusions demonstrate that the cellular balance between NIPBL and PDS5 abundance sets the overall rate of cohesin extrusion, advancing a quantitative mechanistic model that frames the cellular significance of the previously proposed molecular competition model (45, 46) and the observation that NIPBL binding to cohesin is short-lived (*39*, *79*). Collectively, the emerging scenario is that NIPBL binding to chromatin-bound cohesin would enable extrusion; when NIPBL unbinds, extrusion would pause until the next binding event. PDS5 would shield cohesin from NIPBL, thereby extending the interval between extrusion bursts (Fig 6C). A key prediction from this hypothesis is that increasing NIPBL expression in cells with normal PDS5 levels should accelerate extrusion. Indeed, we observed that simply overexpressing NIPBL induces vermicelli formation (fig. 6D), consistent with faster extrusion rates.

Finally, given the haploinsufficient nature of NIPBL for human development, we sought to understand how the rate of cohesin loop extrusion scales with NIPBL expression levels. For this we generated a mouse ESC line expressing NIPBL-FKBP fused to GFP, alongside RAD21-Halotag. This allowed precise monitoring of intermediate NIPBL levels through flow cytometry, after titrating its expression level to generate a range of partial dosage reductions that include and extend beyond the expression levels observed in Cornelia de Lange syndrome. Consistent with the original NIPBL-FKBP line (lacking GFP) (14), basal NIPBL levels were slightly reduced (∼80% of WT untagged cells) as estimated by Western blot, likely reflecting slight leakiness of the degron system (fig. S1A-B). Using limiting amounts of dTAG we generated NIPBL-FKBP-GFP cells with ∼50% and ∼22% NIPBL levels (fig. S12B) and profiled them by Hi-C, and analyzed them side-by side with previously described NIPBL-FKBP undepleted (80% NIPBL levels) and depleted to ∼7%. We measured bound RAD21-Halotag levels by flow cytometry (fig. S12C), enabling calibration of simulated extruder separation. We performed a series of polymer simulations sweeping across a range of extrusion rate multipliers to identify which rate values best fit experimental Hi-C *P(s),* for each NIPBL dose (fig. 6E). We found that inferred rates increased with NIPBL levels across the five samples (fig. 6E). These analyses indicate that NIPBL dosage quantitatively tunes the rate of DNA loop extrusion in cells, thereby modulating genome folding, with immediate implications for clinical contexts where dosage is only partially reduced, such as Cornelia de Lange syndrome.

## Discussion

### Connecting the dynamics of individual cohesin molecules to chromosome-scale organization

While prior genetic and *in vitro* studies identified foundational roles for cohesin cofactors, they offered primarily qualitative accounts of how these factors relate to loop extrusion and control genome organization in cells. In contrast, the quantitative framework we establish here connects cofactor dosage to distinct kinetic parameters of extrusion, accounting for how extrusion dynamics govern chromosome morphogenesis across scales. Our models readily explain observations that had remained purely phenomenological - such as how co-depleting NIPBL and WAPL leaves genome architecture largely intact despite dramatically altering cohesin dynamics - by showing how compensatory shifts in extrusion rate and cohesin lifetime preserve processivity. More broadly, these findings offer a long-sought mechanistic insight into how modest reductions in NIPBL expression, such as those found in Cornelia de Lange syndrome, can disrupt genome organization even when overall cohesin levels on chromatin remain high (*80*, *81*).

A key advance in our computational approach was to explicitly model extrusion rate for comparison with experimental datasets. This enabled us to infer a prominent role of NIPBL abundance on extrusion rate, beyond its effect on loading. Our estimated rate of ∼0.25 kb/s in wild-type ESCs reflects the average overall chromatin-bound cohesin molecules, only a subset of which is expected to be associated with NIPBL and actively extruding at any given time (*39*). Indeed, high-resolution live-cell tracking of DNA loci has reported rare bursts reaching ∼2.7 kb/s, which might specifically correspond to the rate of NIPBL-bound, actively extruding cohesin (*82*). We speculate that variations in the abundance of NIPBL and chromatin-bound cohesin across cell types could account for differences in estimated rates across studies (e.g., 0.85 kb/s in HeLa (*83*)).

Our simulations and experimental titrations also revealed the prevalence of cohesin-cohesin collisions, even in the basal regime. Indeed, accounting for reduced collisions was essential for accurately predicting *P(s)* changes after partial RAD21 depletion. Still, our observations call for caution when interpreting derivatives of *P(s),* as the quantitative relationship with loop size is incompletely determined in biologically relevant scenarios (*68*). Indeed, in simulations, the inflection point in *P(s)* was substantially less than loop size, which in turn was substantially less than the processivity of an unimpeded extruder (fig. S5C). While frequent cohesin collisions can lead to stacking of loops, higher-order rosettes are however rare in cells (*52*) and likely transient because of WAPL-mediated unloading. These observations for interphase cohesin parallel recent observations in mitosis, where condensin-condensin encounters mainly result in collisions rather than bypass (*84*).

### Reframing the roles of NIPBL in cohesin dynamics

Reconstitution studies demonstrated that the association of NIPBL with cohesin is continuously required for extrusion *in vitro* (*40*, *41*, *79*). This led to the proposal that, beyond its effect on cohesin loading, NIPBL functions as a processivity factor during extrusion (*6*), and that there may be no *bona fide* loading complex for extrusive cohesin (*40*). Our data supports a role in cohesin processivity and specifically ascribes a quantitative relationship between cellular NIPBL abundance and the average extrusion rate. Since NIPBL only associates with cohesin intermittently, rapidly binding and unbinding while cohesin is on DNA (*39*, *40*), we speculate that NIPBL depletion slows extrusion rates by increasing the time a chromatin-bound cohesin molecule would spend between two NIPBL association events.

In terms of the apparent contribution of NIPBL to cohesin loading, the observation that NIPBL depletion reduces cohesin levels on chromatin does not necessarily imply that NIPBL performs a dedicated loading reaction directly onto DNA. Indeed, NIPBL may license extrusion after cohesin associates with DNA, potentially via the STAG1/2 subunit (*11*, *85–87*). In this context NIPBL would still effectively act as a loader, by enabling cohesin to engage in the extrusion cycles that prevents its dissociation from chromatin until WAPL-mediated release (*83*). Future work will clarify to what extent this relates to the different requirements of cohesin-STAG1 and cohesin-STAG2 on NIPBL (*11*).

### Dual roles of cellular PDS5: lowering extrusion rates and promoting unloading

We disentangle contributions of PDS5 on extrusion rate from unloading. The molecular basis of the competition between NIPBL and PDS5 remains incompletely understood. *In vitro* assays revealed that PDS5 docks to cohesin on the same site as NIPBL, and their binding is mutually exclusive: PDS5-bound cohesin has no ATPase activity, and cannot extrude loops (*40*, *45*, *46*). Yet, it remains unclear whether PDS5 actively displaces NIPBL in cells, or whether it only binds cohesin after NIPBL spontaneously dissociates. Better tools

will be needed to compare the dissociation kinetics of NIPBL and PDS5 *in vivo*, and to examine their interplay with the dynamics of cohesin acetylation, which is known to alter the affinity of PDS5 for cohesin (*74*) and interfere with WAPL unloading (*88*). The exact role of PDS5 during unloading remains to be clarified. Possible requirements for release include: simultaneous PDS5 and WAPL co-binding, sequential binding (*83*, *89*, *90*), or independent pathways operated by PDS5 and WAPL (*7*).

### The interplay of cohesin-cofactors impart exquisite tunability to interphase extrusion

Our findings demonstrate that altering cohesin cofactor levels can selectively influence certain extrusion parameters while buffering others. For example, concomitant downregulation of PDS5 and NIPBL preserves extrusion rates while increasing cohesin lifetime, and downregulation of both WAPL and NIPBL preserves overall processivity despite elevated cohesin lifetime. Notably, while increases in extrusion rate and cohesin lifetime both enhance processivity equally, we demonstrate that they lead to distinct outcomes for genome folding. We hypothesize that this balancing act provides cells with flexible yet precise control over long-range chromosomal interactions. Such modularity may be key to orchestrate communication between *cis*-regulatory elements while independently modulating genome compartmentalization to facilitate long-term epigenetic memory (*91*).

### Developmental relevance

The expression levels of cohesin and its regulators vary across cell types, providing regulatory opportunities to fine-tune specific aspects of genome folding and achieve distinct genomic functions. For example, WAPL, PDS5A, and PDS5B levels are especially high in ESCs, where extrusion processivity appears lower (*92*), and their expression is crucial for the maintenance of pluripotency and proper silencing of facultative heterochromatin (*23*, *53*, *93*, *94*). In contrast, other cell types naturally express very low levels of WAPL and experience higher extrusion processivity, as seen in pro-B cells, where high cohesin processivity facilitates long-range VDJ recombination at the *Igh* locus (*17*, *18*), and in olfactory sensory neurons, where it allows long-range enhancer activation across the large *Pcdh* loci (*19*, *20*). By modulating the chromosomal interactions that underlie topologically associating domains, cell type-specific changes in extrusion kinetics offer a dynamic mechanism to reconfigure regulatory landscapes across cell types.

Loop extrusion is a major player in the morphogenesis of meiotic chromosomes across species (*95*, *96*). Consistent with the role of PDS5 in lowering extrusion rates, disabling PDS5 in germ cells shortens the meiotic chromosome axis and affects crossover frequency in yeast and mice (*29*, *97*). It is tempting to speculate that, akin to our observations in interphase, the NIPBL:PDS5 ratio directly regulates extrusion rates in meiotic cells, with immediate relevance for the evolution of genetic variation.

### Insight for cohesinopathies

Our work underscores that NIPBL and cohesin abundance differentially influence chromosome folding, bringing new perspectives to the understanding of cohesinopathies. Downregulation of cohesin itself results in fewer but longer DNA loops, as extruders collide less often. In contrast, reduced NIPBL abundance slows the growth of DNA loops, and can cause stronger increases to compartmentalization than cohesin downregulation alone. These mechanistic insights shed light on how altered cohesin or NIPBL expression lead to distinct genome folding defects and are associated with divergent disease outcomes (*98*). Recognizing that NIPBL

expression quantitatively tunes extrusion rates is especially relevant to our understanding of the molecular etiology of Cornelia de Lange syndrome: lower extrusion rates not only shorten DNA loops but also alter their dynamics, dysregulating transcription of gene loci in cell types where they rely on distal enhancers, ultimately derailing developmental processes (*14*). A lower rate of extrusion can account for the impaired distribution of cohesin along chromosomes reported in models of Cornelia de Lange syndrome (*16*, *81*). These considerations also apply to NIPBL mutations that lower the extrusion activity of cohesin without necessarily altering NIPBL abundance directly (*99*), and to mutations in genes other than NIPBL that alter its availability or its stoichiometry to other cohesin cofactors – as observed in Ewing sarcoma (*100*).

Transcriptional and cell differentiation defects caused by NIPBL impairment can be prevented or even reversed through inhibition of WAPL (*47*, *49*, *69*, *101*), highlighting significant therapeutic potential. By revealing that this compensation hinges on the rate of loop extrusion rather than cohesin loading, our findings will inform rational approaches to therapeutic intervention in Cornelia de Lange syndrome and other cohesinopathies.

### Summary

Altogether, this work illuminates how the molecular abundance of cohesin cofactors regulate different steps of the extrusion process, and how their tunable expression can modulate multiple aspects of higher-order genome organization. This loop extrusion control system, while operating at the molecular level, enables the emergence of higher-order structural and functional properties of the genome. Our findings thus opens new systems-level avenues for approaching the molecular processes that underlie genome function and stability in health and disease.

### Limitations of the study

As our models are calibrated for extruder separation using experimental data, the accuracy of our extrusion parameter estimates is constrained by how precisely extrusive chromatin-bound cohesin can be quantified. Future developments in quantification methodologies will sharpen the estimates of loop extrusion parameters derived with our framework. Additionally, cohesin extrusion parameters may vary across the genome, which genome-wide *P(s)*-based analyses do not capture. Our baseline model was calibrated to published estimates of absolute cohesin abundance (from FCS and mass spectrometry) and cohesin lifetime (from FRAP), which might be imprecise. However, such uncertainty in baseline parameters is not expected to affect their inferred fold changes after cofactor perturbations. Our simulations do not model extrusion barriers such as CTCF, the transcription machinery or MCM complexes. As these factors have minor effects on genome-wide Hi-C *P(s)* (*7*, *102–104*), they are not expected to significantly alter our parameter inferences. Another limitation is the use of cycling cells, due to the technical challenge of arresting large numbers of ESCs and releasing them while maintaining viability, which precluded bulk assays in precisely staged populations. Finally, while degron-mediated strategies are useful to circumvent the essentiality of cohesin cofactors, the depletions are not complete, and basal expression is often lower than in untagged cells (especially for AID tags).

## Supporting information

Supp_Table_1_Cell-Lines-Vectors-sgRNAs_v1.xlsx

Supp_Table_2_HiC_mapping_statistics.xlsx

vidS1.mp4 - Differential effects of chromatin lifetime and extrusion rate on single-cohesin behavior

vidS2.mp4 - Simulated impacts of extrusion parameters on local chromatin structure and dynamics

Annotated plasmid maps.zip

## Acknowledgements

We thank Elzo de Wit for helpful discussions and sharing WAPL-AID ESCs as well Jan-Michael Peters, Gordana Wutz, Leonid Mirny and Edward Banigan for communicating unpublished results. We are grateful to Daniele Canzio, Benoit Bruneau and the Nora lab for critical reading of the manuscript. We are deeply appreciative of the Luke Lavis lab and the Open Chemistry team (Janelia) for the gift of JF and JFX dyes. We are thankful to: Kevin So in the lab of Benoit Bruneau for providing technical support isolating PDS5-AID cells at the early phase of this project, Pia Mach for advice optimizing microscopy analyses of ESCs, Laurent Gelman and the Facility for Advanced Imaging and Microscopy (FAIM) at the FMI. We thank the Nora and Fudenberg labs for stimulating discussions.

## Funding

Grant NIH 1R35GM142792-01 to EN, the Chan-Zuckerberg Biohub San Francisco Investigator program to EN, the Hellman Foundation UCSF Fellows program to EN. Research in GF’s lab is supported by NIH R35GM143116 to GF. Research in LG’s lab is supported from the Novartis Research Foundation, the Swiss National Science Foundation (grant no. 310030_192642). KH and DW were supported by National Institute of Health training grant 5T32HD007470 for the UCSF Developmental and Stem Cell Biology Graduate Program and the UCSF Discovery Fellows Program, and KH received support from the California Institute for Regenerative Medicine Scholars Training Program education grant EDUC4-12812 for UCSF. DW was supported by the National Science Foundation GRFP and NICHD Predoctoral Training in Developmental Biology Grant 2T32HD007470. EA was supported by postdoctoral fellowships from the Helen Hay Whitney Foundation and the Regenerative Medicine Scholars Training Program education grant EDUC4-12812 for UCSF. Part of the sequencing was performed at the UCSF CAT, supported by UCSF PBBR, RRP IMIA, and NIH 1S10OD028511-01 grants.

### Author contributions

RS performed tissue culture, genome editing, fractionation-flow-cytometry, Hi-C, RNA-seq, Western blot and contributed to RNA-seq analyses. MMCT performed polymer simulations and analysis of experimental Hi-C data. NL performed iFRAP experiments and analyses under the supervision of LG. HR contributed to polymer simulations. EA contributed to genome editing experiments. KH and DW contributed to the RNA-seq analyses. GF developed preliminary polymer simulations and supervised polymer simulations. EPN performed most genome editing experiments, fixed cell microscopy, cell synchronization experiments, supervised all wet lab experiments except iFRAP, analyzed experimental Hi-C, RNA-seq and fixed cell microscopy. EPN and GF jointly designed the project. EPN and GF wrote the manuscript and assembled the figures with input from other authors.

### Data and material availability

Cell lines and plasmids can be made available from the corresponding author upon request and MTA. Hi-C and RNA-seq data have been deposited to the Gene Expression Omnibus (GEO) database: GSE279296 token odqhwcaetvczhwh for Hi-C and GSE305208 token algbqcocxxwhril for RNA-seq. Requests related to polymer simulations will be fulfilled by co-corresponding author Geoffrey Fudenberg. Microscopy data reported in this paper will be shared by the lead contact upon request. Any additional information required to reanalyze the data reported in this paper is available from the lead contact upon request.

### Supplementary Materials

Figs. S1 to S12

Tables S1 to S2

Videos S1 to S2

Other file type S1

## Methods

### Plasmid Construction

Plasmids were assembled using Gibson assembly (SBI MC010B-1) or restriction-ligation (NEB Quick Ligase M2200). The list of plasmids generated in this study can be found in the supplementary table and their annotated sequence maps as supplementary information.

### Culturing mouse embryonic stem cells (ESCs)

ESCs were cultured in DMEM+Glutamax (ThermoFisher 10569044) supplemented with 15% Fetal Bovine Serum (ThermoFisher 10437028), 550µM b-mercaptoethanol (ThermoFisher 21985-023), 1X non-essential amino-acids (ThermoFisher 11140050), 10^4^ U/mL of Leukemia inhibitory factor (Millipore ESG1107), PD0325901 (APExBIO A3013), CHIR-99021 (APExBIO A3011). Cells were maintained at a density of 0.2-1.5x10^5^ cells/cm2 by passaging using TrypLE (ThermoFisher 12605010) every 24-48h on 0.1% gelatin-coated dishes at 37°C and 5% CO_2_. 0.1% gelatin was prepared in 1X PBS using Gelatin powder (Sigma Aldrich G1890). Medium was changed daily when cells were not passaged. Cells were checked for mycoplasma infection and tested negative.

### Degron inductions

Degradation of proteins tagged with the Auxin-Inducible Degron (44 amino-acid AID* truncation with residues 71–114 (Nora et al., 2017)) was induced by adding the auxin analog Indole-3-acetic acid sodium (IAA, Sigma Cat #I5148-2G) to a final concentration of 500μM (from a 1000x filter-sterilized stock diluted in water). Degradation of proteins tagged with the FKBP degron (FKBP12^F36V^ (Nabet et al., 2018)) was induced by adding dTAG-13 (abbreviated dTAG throughout the manuscript - Sigma Cat, SML2601) to a final concentration of 500nM (from a 1000x stock diluted in DMSO). We noticed equivalent degradation efficiency in ESCs compared to using the dTAG-v1 inducer at 500nM.

### Genome editing of ESCs

For transfection, plasmids were prepared using the Nucleobond Midi kit (Macherey Nagel 740410.5) followed by isopropanol precipitation. Constructs were not linearized. Transfections were typically done using the Neon system (Thermo Fisher) with a 100 µL tip and 1 million cells at 1400 V, 10 ms, 3 pulses. To create knock-in cells, ESCs were co-transfected with 5 µg of Cas9-sgRNA expressing vector and 10-15 µg of targeting construct.

sgRNAs were cloned either into a Cas9 vector (pX330, Addgene #42230) or Cas9-2A-puro vector (pEN243, identical to pX459 Addgene 62988). Homology arms of targeting vectors ranged from 800 base pairs (bp) - 1 kilobase (kb) in length.

After electroporation, cells were seeded in a 9 cm^2^ well and left to recover for 48 hours. Cells were plated at limited dilution and grown for around 8 days changing medium every other day until single colonies could be picked. Individual colonies were genotyped by polymerase chain reaction (PCR) and validated with Sanger sequencing. Homozygous or heterozygous clones were identified, expanded, validated by western blot, and cryopreserved. For serial editing, resulting validated clonal lines were used to repeat the transfection process with another targeting construct.

When the targeting construct contained an FRT flanked resistance cassette, we adopted the following strategy to avoid having to subclone the cells again after removal of the resistance cassette: after transfection of the targeting construct containing the FRT flanked resistance cassette, we added antibiotic in the medium 48h after transfection and kept the cells under selection for at least 5 days. Final antibiotic concentrations: Puromycin: 1 µg/mL; Blasticidin: 10 µg/mL; Hygromycin: 200 µg/mL; Geneticin: 200 µg/mL. We then transfected this heterogenous pool with pCAGGS-FlpO using the Neon electroporation system with a 10 µL tip and 0.1 million cells at 1400 V, 10 ms, 3 pulses, using 0.5µg DNA. Cells were seeded in a 9 cm^2^ well and subcloned without antibiotic after 48h. PCR genotyping and antibiotic resistance tests identified homozygous knockin clones with the FRT-flanked selection cassette removed.

The list of cell lines, genotypes, order of transfection and corresponding vectors is provided as a supplementary table.

### Western blotting

ESCs were dissociated using TrypLE, resuspended in culture medium, pelleted at 100*g* for 5 minutes at room temperature. Cell pellets were washed in PBS, pelleted again, snap frozen in dry ice and stored in −70°C freezer. In total, 10-20 million cells were used to prepare nuclear extracts. Cell pellets stored at −70°C were taken out on ice, resuspended in 10 mM HEPES (pH 7.9), 2.5 mM MgCl_2_, 0.25 M sucrose, 0.1% NP40, 1 mM DTT, and 1X HALT protease inhibitors (ThermoFisher 78437) and kept on ice for 10 minutes to allow swelling. After centrifugation at 500*g* at 4°C for 5 minutes, the supernatant was discarded and the pellets were resuspended on ice in 25 mM HEPES (pH 7.9), 1.5 mM MgCl_2_, 700 mM NaCl, 0.5 mM DTT, 0.1 mM EDTA, 20% glycerol, 1 mM DTT, and 250 U benzonase (Sigma-Aldrich E1014), and incubated on ice for 30-60 minutes. The nuclear lysates were centrifuged at 18,000*g*, 4°C, 10 minutes, to eliminate the cellular debris. Protein concentration from the supernatants was measured using Pierce BCA Kit (ThermoFisher 23225).

Equal amounts of protein lysates were heated in 1X Laemmli buffer (Biorad 1610747) at 95°C for 10 minutes and subjected to electrophoresis on a 4-15% polyacrylamide gel (Biorad 17000927). For NIPBL westerns, 60 µg protein was loaded. The separated proteins were transferred onto a polyvinylidene difluoride (PVDF) membrane (Millipore IPFL00010) using phosphate-based transfer buffer (10 mM sodium phosphate monobasic, 10 mM sodium phosphate dibasic) at 4°C, 600 mA, 2 hours. After the completion of transfer, membranes were blocked in the 5% blocking buffer (5% skimmed milk prepared in 1X TBS-T) and incubated overnight at 4°C with the primary antibodies prepared in the 5% blocking buffer. Next day, the membranes were washed four times with 1X TBS-T buffer (20 mM Tris-Cl buffer-pH 7.4, 500 mM NaCl and 0.1% Tween 20) and incubated with the appropriate secondary antibodies conjugated with horseradish peroxidase (HRP) for an hour at RT (typically 1:5000). Following this, the membranes were again washed four times with 1X TBS-T buffer. The blots were developed using Pierce ECL western blotting substrate (ThermoScientific 32209) and detected using Chemidoc XRS+ imager (Biorad).

For validating homozygous knockin and degron depletion efficiency we used the following antibodies: PDS5A: Bethyl Labs A300-089A 1:1000 PDS5B: Bethyl Labs IHC-00381 1:1000 TBP (loading control): Cell Signaling technology 8515S 1:2000 RAD21: Millipore 05-908 1:1000 NIPBL: sc-374625 1:1000 MAU2: Abcam ab183033 1:1000

### Flow cytometry to measure total and chromatin bound RAD21

ESCs were seeded at a density of 0.5 x 10^6^ per well of a gelatinized 6-well plate. Next day, the degron containing ESC lines were treated with either IAA or dTAG-13 or both for 6 hours. At the end of 6 hours, media was aspirated out and replaced with medium containing 100nM or 500nM of Halo-ligand JFX549 or JFX646 (kind gift of Luke Lavis, HMMI Janelia Research Campus) and specified concentration of degron inducers (IAA and/or dTAG-13). The cells were incubated at 37°C, 5% CO_2_ for 30 minutes to allow JFX549 to bind the Halo-tag on RAD21. After 30 minutes, media was aspirated, cells were washed twice with 1X PBS and fresh media containing the specified concentration of degron inducers (IAA and/or dTAG-13) was added to the well. Cells were incubated at 37°C, 5% CO_2_ for 30 minutes to allow for unbound JFX549 or JFX646 to diffuse out of the cells. Following 30 minutes incubation, cells were rinsed with 1X PBS, dissociated using TrypLE, resuspended in fresh culture medium containing appropriate amounts of IAA or dTAG-13 and centrifuged at 100g for 5 minutes at RT. Supernatant was aspirated and the cell pellets were resuspended in 1 ml of 1X PBS. Resuspended cells were equally divided into two tubes to measure total RAD21 and bound RAD21 and spun at 100g for 5 minutes at RT. Supernatant was discarded.

For measuring total RAD21, cell pellets were resuspended in 100ul of 4% formaldehyde (prepared in 1X PBS, diluted from 16% methanol-free formaldehyde - ThermoFisher 28908) and incubated at RT for 15 minutes. After incubation, 1 ml of ice-cold 1% BSA (diluted from 3% BSA in 1X PBS) was added and cells were spun at 300g for 5 minutes. Supernatant was discarded. Cell pellets were resuspended in 100ul of ice-cold CSK-T buffer (25mM HEPES pH7.4, 150mM NaCl, 1mM EDTA, 3mM MgCl2, 300mM Sucrose, 0.5% TritonX-100) on ice and incubated on ice for 10 minutes. After incubation, 1 ml of ice-cold 1% BSA (diluted from 3% BSA in 1X PBS) was added and cells were spun at 300g for 5 minutes. Supernatant was discarded. Cell pellets were resuspended in 300ul of 1mg/ml DAPI solution prepared in 1X PBS. Incubated for 5-10 minutes and acquired using BD Versa flow cytometer using PE-A (for JFX549), APC-A (for JFX646) and V450 (for DAPI) channels. 100,000 events were acquired.

For measuring chromatin-bound RAD21, cell pellets were resuspended in 100ul of ice-cold CSK-T buffer (25mM HEPES pH7.4, 150mM NaCl, 1mM EDTA, 3mM MgCl2, 300mM Sucrose, 0.5% TritonX-100) on ice and incubated on ice for 10 minutes. After incubation, 1 ml of ice-cold 1% BSA (diluted from 3% BSA in 1X PBS) was added and cells were spun at 300g for 5 minutes. Supernatant was discarded, pellets were resuspended in 100ul of 4% formaldehyde (prepared in 1X PBS, diluted from 16% methanol-free formaldehyde) and incubated at RT for 15 minutes. After incubation, 1 ml of ice-cold 1% BSA (diluted from 3% BSA in 1X PBS) was added and cells were spun at 300g for 5 minutes. Supernatant was discarded and pellets were resuspended in 300ul of 1mg/ml DAPI solution prepared in 1X PBS . Incubated for 5-10 minutes and acquired using BD Versa flow cytometer using PE-A (for JFX549) and V450 (for DAPI) channels. 100,000 events were acquired.

For complete depletion of PDS5A/B or WAPL, respective degron lines were treated with 500 μM of IAA for 6 hours. For complete depletion of NIPBL, the NIPBL-FKBP degron line was treated with 500nM of dTAG-13 for 6 hours. For co-depletion of PDS5A/B and NIPBL, WAPL and NIPBL, respective degron lines were treated with 500 μMof IAA and 500nM of dTAG-13 for 6 hours. To achieve partial RAD21 depletion in the background of complete PDS5A/B-depletion, cells were treated with 500 μMof IAA (to completely deplete PDS5A/B) and 10nM or 25nM dTAG-13 (to partially deplete RAD21) for 6 hours. To achieve partial RAD21 depletion in the background of complete WAPL-depletion, cells were treated with 500 μMof IAA (to completely deplete WAPL) and 8nM or 25nM dTAG-13 (to partially deplete RAD21) for 6 hours. To achieve partial depletion of RAD21, cells were treated with 20nM or 45nM of dTAG-13 for 6 hours.

ESC line devoid of any degron-tag and without Halo-tag on RAD21 served as a negative control. ESC line without any degron-tag and with Halo-tag on RAD21 served as a positive control (no degron). The median fluorescence for PE-A channel from the single cell gated for total RAD21 and bound RAD21 was subtracted from the respective values coming from the cells devoid of Halo-tag. This eliminated any non-specific background from the Halo ligand. Mean fluorescence for RAD21 was plotted for both total RAD21 and bound RAD21 for No degron (RAD21^Halo^) and different degron lines.

Percent bound RAD21 was calculated using following formula, taking the mean fluorescence across each cell population:

Step 1: For all samples, remove aspecific signal by subtracting fluorescence levels of Halo-labeled untagged cells (not expressing the halotag) from the fluorescence levels of Halo-labeled RAD21-Halotag cells

Step 2: Calculate % Bound RAD21 as follows:

i. %Bound RAD21 = (RAD21 after pre-extraction)*100/(Total RAD21)
ii. Percent bound RAD21 relative to No degron was derived as follows:

%Bound RAD21 relative to No degron = (Bound RAD21 in degron line)*100/(Bound RAD21 in No degron)

### Cell cycle analysis by EdU incorporation and DAPI staining

The Click-iT Plus EdU Alexa Fluor 647 Flow Cytometry Assay Kit (Thermo Fisher Scientific C10634) was used following the manufacturer’s instructions. We used 20 µM EdU, pulsed for 1 hour and 30 minutes, and proceeded with 1 million cells, incorporating DAPI after EdU staining.

### Mitotic synchronization and release, cell cycle analysis by DAPI and RAD21 vermicelli microscopy

Three T75 flasks per genotype were incubated overnight (9h) with 100 ng/mL Nocodazole (from a 1/40 dilution followed by a 1/500 dilution from a 2mg/ml stock, Sigma-Alrdich M1404). IAA (auxin analog) was then added at 500 μM final to the culture medium to induce WAPL or PDS5AB depletion for 3h. Mitotic shake-off was then performed by firmly tapping the flaks, and the supernatant culture medium containing the dislodged mitotic cells was collected, pooling the three T75 flasks in a 50mL conical tube. Cells were spun, resuspended in 25mL 1XPBS containing IAA, spun again, resuspended in 25mL 1XPBS containing IAA again counted, spun once more and resuspended at 1 million cells per mL in ES culture medium with nocodazole but with 500 μM IAA. This procedure yielded around 10M cells across the three T75 flasks. The ‘0h release’ timepoint was immediately collected by aliquoting 1M cells. For the other timepoints 1M cells were seeded in a gelatinized tissue culture 9 cm^2^ well for later hourly collection, in ESC medium with 500 μM IAA. For the first two hours cells typically only loosely attached and could be singularized by pipetting. For later timepoints cells were detached and singularized as during regular passaging using TryplE. For each time-point we collected three fractions. The synchronized/released cell pellet was resuspended in 1XPBS with 500 μM IAA and split in two fractions:

- Microscopy fraction (to quantify RAD21 vermicelli scores): 0.1M to 0.2M cells were seeded in a well of a glass-bottom 96-well plate (Greiner 655892). To ensure cells adhere the glass had been pre-coated with Poly-L Lysine. To coat the glass, we incubated 50 μL per well of 0.01 % (w/v) Poly-L-lysine solution (P8920-100ML) diluted in water for 10 minutes at room temperature and dried for at least 30min (up to days). After 10 minutes of incubating the cells onto the coated glass excess volume was removed by pipetting (adhered cells stayed on the coated glass a monolayer of single-cells) and cells were incubated with 100uL of 3% formaldehyde diluted in 1XPBS (from a 37% solution Electron Microscopy Sciences 15686). After 10 minutes wells were rinsed twice with PBS and left in 100uL 1xPBS until all the timepoints were collected. Fixed cells were kept at least overnight at 4°C once all timepoints were collected. For Halotag ligand reaction and H3S10Ph immunostaining, cells were first permeabilized by incubating in 0.5% Triton in 1xPBS for 5 minutes, and rinsed three times with 1xPBS. Then cells were blocked by incubating with 1%BSA in 1xPBS 15 minutes. Incubation was changed for a 1/5000 dilution of anti-H3S10Ph antibody (Millipore 06-570) in 1%BSA 1xPBS for 30min, followed by three 5-minute washes in 1xPBS. Cells were then incubated with a 1/10000 of goat anti-rabbit Alexa-647 (ThermoFisher 21245) and 100nM JFX549 for 30 minutes at room temperature. After three 5-minute washes in 1xPBS, cells were incubated with 1μg/mL DAPI in 1xPBS 10 minutes, rinsed twice with 1xPBS and left in 1xPBS until imaging, within 24h. Imaging was performed on a spinning disk microscope (Zeiss) with a 60x 1.4NA oil objective equipped with 405nm, 488nm, 561nm, 640nm excitation lasers at the Histology and Light Microscopy Core of the Gladstone Institutes. Cells were imaged in 3D using a step size of 0.3μm.

- Flow cytometry fraction (to assess overall synchronization and release efficiency): 400 μL of 1XPBS was added to a 100μL aliquot of cells and transferred into a FACS tube. 70% ethanol was then added dropwise while gently vortexing, and tubes were stored at least overnight at -20. Flow-cytometry fractions for each time-point were transferred to a 1.5mL tube, spun, resuspended in 1mL 1XPBS, spun again, resuspended in 1mL 1XPBS with 0.5% Triton X-100 1μg/mL DAPI and incubated at room temperature 30min. Cells were then processed on a BD Versa flow cytometer using V450 (for DAPI) channels.

### RAD21 vermicelli microscopy from asynchronous ESCs

For RAD21-Halotag imaging in asynchronous cells, adherent cultures were obtained by seeding 100,000 ESCs per well of a laminin-coated glass-bottom 8-well slides (Ibidi 80807) for overnight growth, adding IAA for the time indicated in figure panels. To coat the slides, laminin (L2020-1MG) was diluted 1:100 in 1XPBS (10-20µg/ml working solution), and 100-150μL was added per well for at least 2h up to overnight at 37°C (in tissue culture incubator). Laminin was then aspirated from the well prior to dispensing cells, without rinsing. We found laminin coating minimized the size of the ESC colonies, helping with high-magnification imaging.

When ready for staining, adherent cells were washed with 150μL 1XPBS per well, followed by 3% formaldehyde diluted in 1XPBS (from a 37% solution Electron Microscopy Sciences 15686) for 10 minutes. Cells were then rinsed three times with 150μL 1XPBS, permeabilized by incubating in 0.5% Triton in 1xPBS for 5 minutes, and rinsed three times with 1xPBS. Cells were incubated with 100nM JFX549 1μg/mL DAPI in 1xPBS for 30 minutes, washed twice with 1xPBS for 10 minutes and left in 1xPBS until imaging, within 24h. As most asynchronous ESCs are in S/G2 phases H3S10Ph staining was used in this analysis.

Slides were imaged on a spinning disk microscope (Nikon) with a 60x 1.4NA oil objective equipped with 405nm, 491nm, 561nm, 640nm excitation lasers. at the USCF Center for Advance Light Microscopy. Cells were imaged in 3D using a step size of 0.3μm.

### Quantification of RAD21 vermicelli scores from microscopy images

Proprietary file formats (.czi for Zeiss instrument, mitotic release timecourse; .nd2 for Nikon instrument for asynchronous cells) were open in imageJ using Bio-Formats importer(*105*). A series of custom scripts were used first to extract and sort stacks. A mask for nuclei was then created by creating a Z projection and applying a Gaussian Blur (sigma = 2) followed by a Mean filter (radius=5). Filtered nuclei were then manually thresholded for each image and binarized. Touching nuclei were separated using Watershed processing (tolerance = 1.5) and resulting particles were filtered for size (100–450) and circularity (0.6-1) the resulting collection of region or interests was then used to analyze the other channels. The red channel (JFX549 staining RAD21-Halotag) was quantified by extracting the mean and the standard deviation for each nucleus. For the analysis of synchronized cells, the paired infra-red channel (H3S10Ph staining) was quantified by extracting the mean signal. After processing all nuclei across several image stacks the RAD21-Halotag vermicelli score, or coefficient of variation (StdDev/Mean) was calculated in R. For the analysis of synchronized cells, unreleased mitotic cells were eliminated from the analysis by removing the nuclei with H3S10Ph by thresholding the infra-red signal in R.

### Inverse Fluorescence Recovery after Photobleaching (iFRAP)

The cells were imaged on an inverted motorized stand Zeiss AxioObserver7 equipped with CSU-W1 Confocal Scanner Unit (Yokogawa) with Dual T2, a 50um pinhole disk unit, 2 sCMOS cameras (Prime 95B, photometrics),a Visitron VS-Homogenizer,an ASI MS2000 X,Y, ZPiezo drive (300um travel range) and controlled with VisiView (version 6.0.0.35) imaging software (Visitron Systems GmbH). Illumination was achieved with 561 Cobolt Jive (Cobolt, 200mW). For FRAP, illumination was done using a 473 nm FRAP laser. A plan Apochromat 40X/1.3 oil objective was used, resulting in a pixel size of 0.275 μm in x,y. A 405/488/561 dichroic was used. The emission filter in the scan head was a 575 nm long pass. Imaging was done in a humidified incubation chamber at 37° C supplied with 8% CO2.

A ROI was defined on the field of view and few cells (∼10) were selected for FRAP. A rectangular ROI was defined for each nuclei encompassing around 80% of the nuclei area at the focal plane. 2D timelapse images were acquired every 30 s for 250 frames, further referred to as movies. Images were taken at 30% max laser power (∼ 3.528mW) to limit bleaching. The focus was kept using a Zeiss definiteFocus 2 system between every frame. After 4 frames, the FRAP sequence was initiated on the selected nuclei at 100% laser power. For FRAP, a laser of 22 um was focused on the focal plane and was shined on each pixel for 300ms. The focusing of the FRAP laser to the imaging focal plane and calibration of the x,y direction of the laser was done before every acquisition using a dedicated well not used for the imaging. The calibration was done by moving the beam to specific locations on the field of view and adjusting its focus to obtain the smallest beam at the imaging focal plane.

The cells were imaged on a µ-Slide 8 Well glass bottom (Ibidi, 80807). Briefly, a day before imaging ∼200.000 cells were split in Gibco™ FluoroBrite™ DMEM (Gibco™) onto the dish. On the day of the imaging 100nM of Halo-JF-549 ligand (Janelia Fluor® NHS ester Tocris lab, 6147) was added for 30 minutes to the imaging medium. Then cells were washed three times with 1X PBS followed by addition of fresh imaging media.

### Image analysis

The movies were saved as ome.tf2 format in the VisiView software. The images were analyzed using a custom GUI python script. We selected 2 circular ROI for both bleached and unbleached regions and updated their position every 5 frames (2.5 minutes). Missing positions were linearly interpolated. The mean intensity in the unbleached (I_unbleached(t)) and bleached (I_bleached(t) regions at every frame are then extracted.

To account for photobleaching we defined for every movie 10 unbleached nuclei and segmented them using Otsu’s intensity thresholding method. To account for cell motion nuclei are linked using maximum overlap as a linking score, nuclei tracks are then built using the most overlapping nuclei. The mean intensity over time (I_photobleaching(t)) is then extracted. Furthermore, to account for the background intensity, another circular ROI was defined in a place on the image without any cells and (I_background) was extracted and averaged over all frames.

For every movie I_photobleaching(t) - I_background is computed and normalized to 1 as the initial difference in order to obtain a photobleaching curve.

Raw I_unbleached(t) and I_bleached(t) are averaged over all the nuclei in a given movie and the difference I_unbleached(t)-I_bleached(t) is computed and normalized by the bleaching to create the iFRAP signal,

iFRAP = I_unbleached(t)-I_bleached(t)/I_photobleaching(t) -I_background

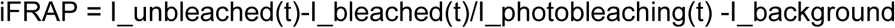

Individual nuclei traces were visually checked and traces corresponding to nuclei leaving the FOV were manually removed. To account for initial cohesin concentration and equilibration of the system after bleaching, the iFRAP signal was further normalized to its level at frame 6 (1.5 minutes after bleaching).

More detailed explanation and raw code is available on https://nesslfy.github.io/cohesin_residence_time/index.html

### RNA-seq

0.5 million ESCs were seeded in a 9 cm^2^ tissue culture well (-/+ dTAG) and collected after 24 hours for RNA extraction. ESCs were dissociated using TrypLE, resuspended in culture medium, spun down at 100g for 5 minutes at RT. Supernatant was discarded and the cell pellets were washed once with 1X PBS. Cell pellets were resuspended in 1 mL of Trizol (ThermoFisher 15596-018), incubated at RT for 5 minutes. 250 μL of chloroform was added, vortexed vigorously, incubated at RT for 10 minutes and spun at 12,000g for 5 minutes at 4°C. Upper aqueous layer was transferred to a fresh tube, mixed with an equal volume of isopropanol, vortexed briefly, incubated at RT for 10 minutes and centrifuged at 12,000g at 4°C for 15 minutes to pellet the RNA. Supernatant was discarded and the pellet was washed twice with 75% ethanol and allowed to air dry for 5 minutes. RNA was resuspended in 50 μL of Nuclease-free water. RNA concentration was measured using Nanodrop and quality was assessed using Bioanalyzer (Total RNA Pico kit- Agilent 5067-1513).

1.5 μg of RNA was treated with the Turbo DNase kit (Invitrogen AM1907), incubated at 37°C for 30 minutes. DNase was inactivated using 2 μL of inactivation reagent by incubating at RT for 5 minutes followed by centrifugation at 13500 rpm for 90 seconds at RT. Supernatant was transferred to a fresh tube. 1000 ng of total RNA was used as input for poly(A) capture and library preparation using the Illumina Stranded mRNA kit (Illumina 20040532) per manufacturer’s instructions. Quality and quantity of libraries were assessed using QuBit and Bioanalyzer (High sensitivity DNA Chip - Agilent 5067-4626). Libraries were sequenced on HiSeq 4000 using 100 bp paired end or NovaSeq 6000 using 150 bp paired end.

### Analysis of RNA-seq data

Reads were trimmed to 100bp paired-end. Alignment of raw RNA-seq data was produced using STAR version 2.7.7a with default parameters and basic two-pass mapping in mm10. GENCODE VM23 (GRCm38.p6) was used as the reference gene set. Size factor normalization and differential expression analyses were done using DESeq2 (version 1.36.0) after filtering out genes with less than 10 raw reads. Principal Component Analysis was conducted in DESeq2 on auxin/dTAG-treated samples with the three replicates separate. Replicates were then collapsed by averaging their PC1 coordinates after PCA, resulting in a single point per condition.

### Hi-C

Hi-C was performed using the kit from Arima Genomics as per manufacturer’s instructions (202103–1577). Briefly, ESCs were dissociated using TrypLE, resuspended in culture medium and counted. 5-10 million cells were spun down at 100g for 5 minutes at RT. Supernatant was discarded, and the cell pellets were resuspended in 5 ml of 1X PBS. Cells were crosslinked using 2% formaldehyde (diluted from 37% formaldehyde - Electron Microscopy Sciences 15686), mixed by inverting the tubes 10 times and incubated at RT for 10 minutes. Quenching was done using the Stop solution 1 provided with the kit, mixed by inverting the tubes 10 times and incubated at RT for 5 minutes. Samples were then incubated on ice for 15 minutes and spun down at 100g for 5 minutes at 4°C. After discarding the supernatant, cells were resuspended in cold 1X PBS, counted, aliquoted at a density of 1x10^6^ cells/tube and spun down at 100g for 5 minutes at 4°C. Supernatant was discarded, cell pellets were snap frozen on dry ice and stored at -70°C.

Fixed cells were thawed and Hi-C was performed as per manufacturer’s guidelines. Libraries were prepared using TruSeq DNA Nano LP kit from Illumina (20015964). Quality and quantity of libraries were assessed using Qubit and Bioanalyzer (High sensitivity DNA Chip- Agilent 5067-4626). Libraries were sequenced on NextSeq 500 using 75 paired end or NextSeq 2000 using 101 bp paired (and then trimmed to 75 before analysis).

### Analysis of Hi-C data

Reads were trimmed to 75bp (if sequenced beyond that) using *cutadapt*(Martin, 2011). Data was mapped to mm10 using the *distiller-nf* pipeline (https://github.com/open2c/distiller-nf), iteratively corrected and converted into .cool format with *cooler*(*106*), only considering read pairs with both sides mapping with high confidence (mapq ≥ 30). After ascertaining the concordance of each replicate using either *cooltools* (Open2C et al., 2024a) or *Genova*(van der Weide et al., 2021) (P(s) curves and aggregate peak analysis), the independent Hi-C replicates (up to two) were merged for each condition. Multiresolution coolers (.mcool) were then generated with *cooler* and balanced using iterative correction. P(s) plots and Hi-C contact maps were generated using *cooltools* (Open2C et al., 2024a). Compartment scores were obtained from the saddle plots using *cooltools* and were normalized to 1 in the limit of full A/B segregation, with a score of 0 denoting ideal A/B mixing. For computations of distance-dependent compartment scores (fig. S7D), experimental observed-over-expected maps were aggregated across all chromosomes in which compartment patterns were found to visually align with the first Hi-C eigenvector (ev1) (chromosomes 1, 2, 4, 5, 6, 8, 9, 11, and 13-19). Hi-C counts were binned against ev1 and partitioned as a function of distance from the main diagonal in 500 kb increments in order to create ensembles of distance-dependent saddle plots, from which compartment scores were similarly calculated.

### Polymer model of ESC chromatin with loop extrusion and compartmentalization

We modeled chr2:66,000,000-116,000,000 using Polychrom-HooMD (https://github.com/open2c/polychrom-hoomd). As previously(*60*), we modeled chromatin as a fiber with monomers of individual diameter 50 nm, each representing 2.5 kb of DNA at a density of 0.2 monomers per unit volume, amounting to a physiological DNA concentration of 0. 01 𝑏𝑝/𝑛𝑚 . Molecular dynamics (MD) simulations were conducted using periodic boundary conditions, and were initiated from a compact conformation which was allowed to expand until the radius of gyration equilibrated over ^7^ MD steps. Based on the comparison of the computed single-locus mean-squared displacements with in vivo measurements in yeast and mammals(*83*), each MD step was estimated to amount to 5 ms of physical time. Genomic positions of A and B compartments were determined in a standard fashion from the first principal eigenvector of the experimental Hi-C correlation matrix obtained in WT embryos at 25 kb resolution, and computed using *cooltools*. While this resolution was sufficient to reproduce the main features of the experimental maps, coarse graining compartment positions to 25 kb contributes to the greater dynamic range of simulated compartment scores. A self-affinity interaction of magnitude 0.05 kT was applied between B-type loci, which was found by iterative optimization to yield a maximum Spearman rank correlation coefficient between the simulated and experimental observed-over-expected maps in wild-type conditions (fig. S2D). To optimize statistical sampling, as well as mitigate potential finite-size effects, each simulation was set up to include 10 identical replicates of the chromosomal region of interest, thus amounting to a total system size of 200,000 monomers (or, equivalently, 500 Mb).

As previously(*60*), we coupled this 3D bead-spring heteropolymer model with a 1D lattice model of loop extruder dynamics. Extruder leg positions from the 1D lattice at each timestep were used to specify positions of additional harmonic bonds between monomers in the 3D simulation, with identical stepping dynamics in A and B compartments. As originally implemented for cohesin extrusion and as recently hypothesized for condensin-condensin encounters(*84*), we assumed that extruders collided without a chance to bypass each-other. Under wild-type conditions, an extruder separation of 185 kb was used to match the loaded cohesin abundance reported in ESCs(*65*). Similarly, the mean extruder residence time on chromatin was set to 1320 s based on FRAP measurements of cohesin dynamics *in vivo* (*56*). During each lattice update in a wild-type 1D simulation, 4000 MD steps were executed. This value was found from a systematic sweep to yield optimal agreement with the experimental contact frequency vs. distance curve in untagged cells, and amounts to an average extrusion rate of 0.25kb/s.

### Simulated Hi-C and contact-vs-distance profiles

*In silico* contact maps were generated from MD simulation trajectories using the monomerResolutionContactMapSubchains function from the contact_maps module as implemented in the polykit library, using the default capture radius (2.3 monomers, 115 nm). Maps were averaged across 10^4^ equilibrated conformations obtained from 10 independent runs. We then determined contact frequencies as a function of genomic distance using the expected_cis function of *cooltools*. For comparisons with experimental data, the obtained *in silico* P(s) curves were linearly interpolated using the interp function of the numpy library to synchronize bin positions to those used for experimental data. *In silico* curves were then rescaled such that the value at 50 kb matches that measured in the *in vivo* sample. Accordingly, the goodness-of-fit parameter (*R*^2^) was defined as

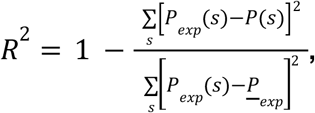

where the summation and average (𝑃_ 𝑒𝑥𝑝) are both performed over the genomic range [50 𝑘𝑏, 5 𝑀𝑏]. For visualization, derivatives of the contact frequency curve (P’(S)) were computed with the `slope_contact_scaling` function from polykit.polymer_analyses.

### *In silico* microscopy

Simulated microscopy images were obtained from individual chromosome conformations by binning computed cohesin spatial positions into discrete voxels of size 100 nm x 100 nm x 100 nm. Optical diffraction effects in the (x, y) plane were mimicked by running the resulting 3D raster through a Gaussian convolution filter of width 250 nm, following which a maximum intensity projection was performed along the z direction. To account for the presence of unloaded cohesins, additional unbound cohesins were placed randomly throughout the simulated volume, with a number determined from FACS estimates for the bound fraction. Vermicelli scores were subsequently defined as in experiments from the ratio of the standard deviation and the mean of the normalized cohesin fluorescence signal.

## Supplementary material

**Fig. S1:**
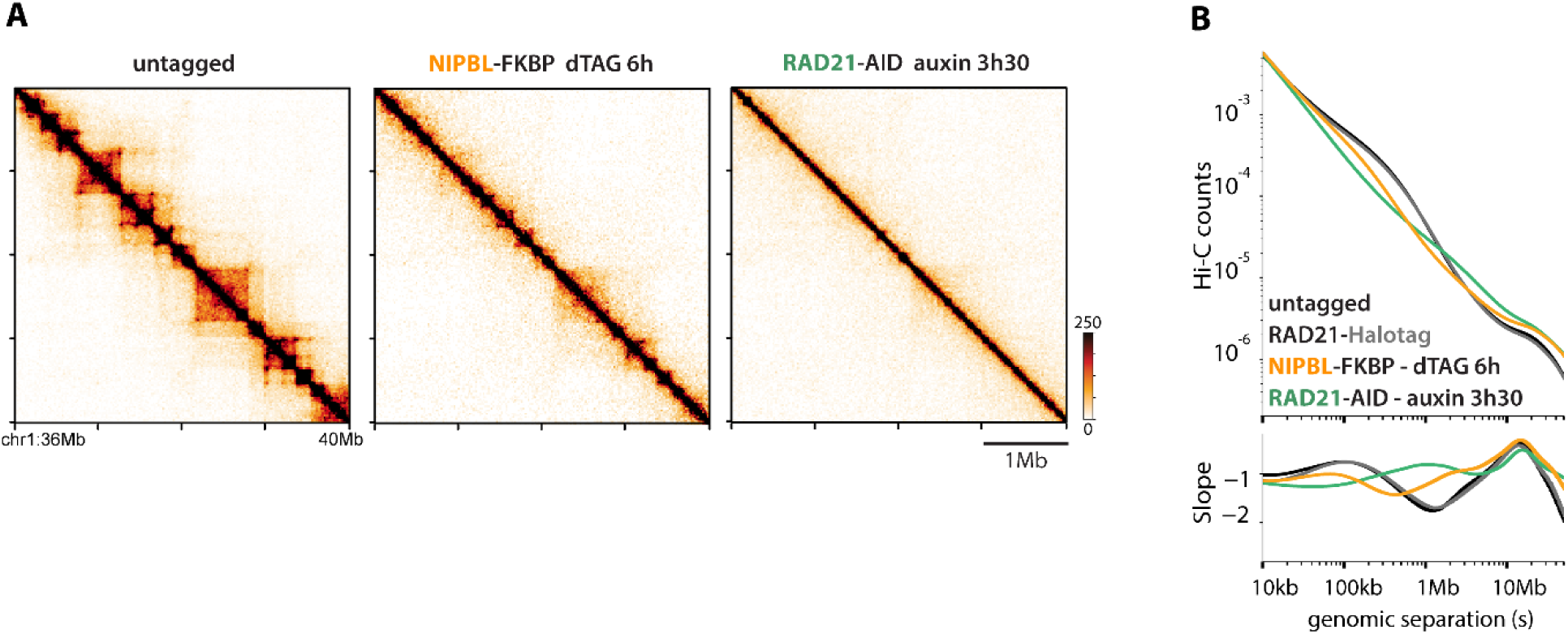
Additional characterization of NIPBL degron cells. (A) Hi-C contact maps (20kb bins) after acute NIPBL depletion (as in Fig.1) or near-complete cohesin (RAD21) depletion with inducible degrons in mouse embryonic stem cells. (B) *P(s)* plots after near-complete degradation of RAD21 or NIPBL and Halo-tagging of RAD21.

**Fig. S2:**
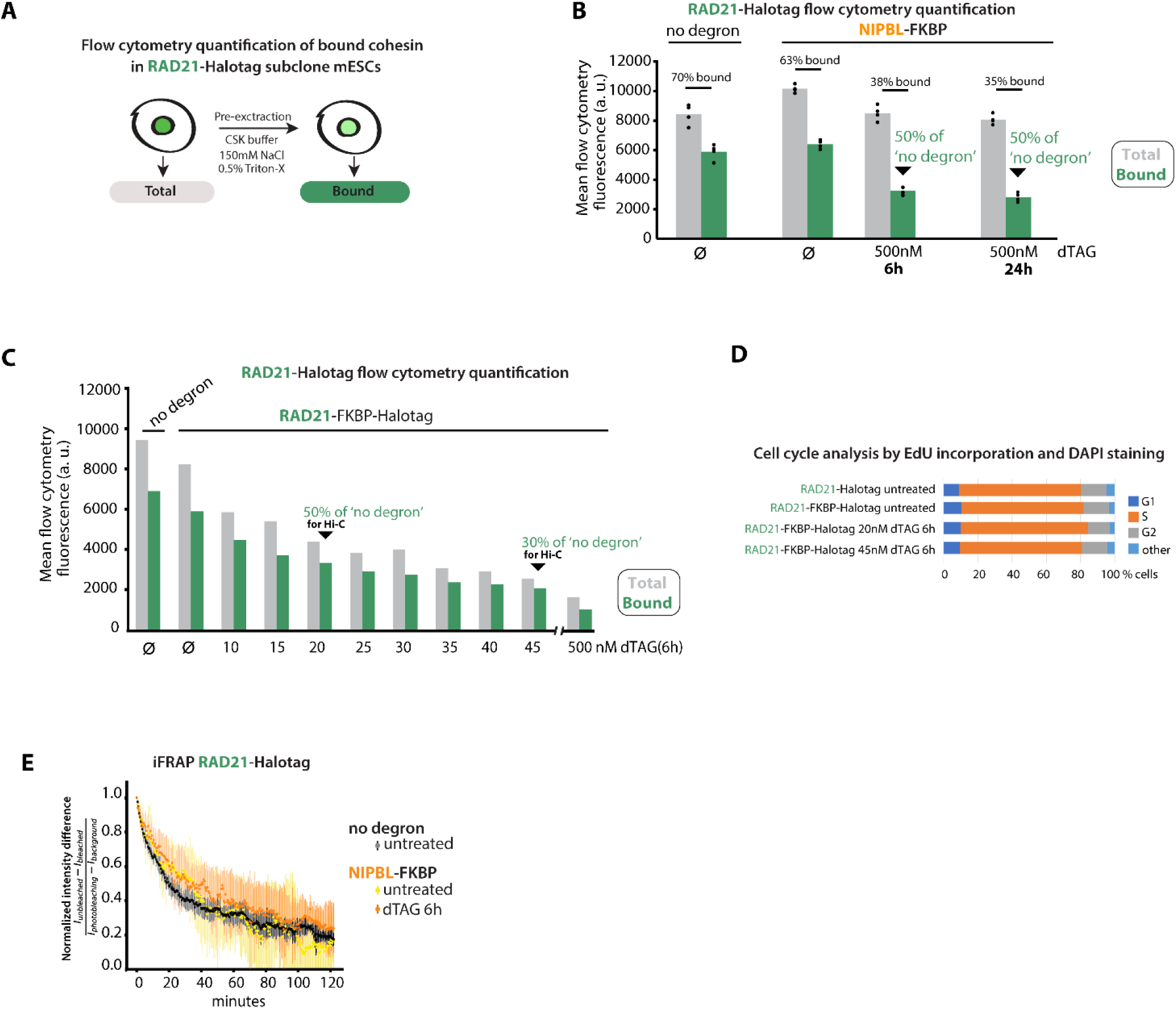
Quantification of total and chromatin-bound cohesin with RAD21-Halotag and iFRAP after NIPBL depletion. (A) Procedure to measure total and chromatin-bound cohesin RAD21 with flow cytometry. (B) Depleting NIPBL for longer than 6h does not further decrease the amount of bound cohesin, even after 24h (1-2 cell divisions), as in Fig. 1. (C) Partial depletions of cohesin levels by dTAG titration in RAD21-FKBP degron cells. (D) Partial cohesin depletion conditions used for Hi-C do not interfere with cell cycle. (E) iFRAP analyses indicate cohesin residence time (lifetime) remains similar after NIPBL depletion. n=16-18 cells. Shown are medians +/- standard deviation.

**Fig. S3:**
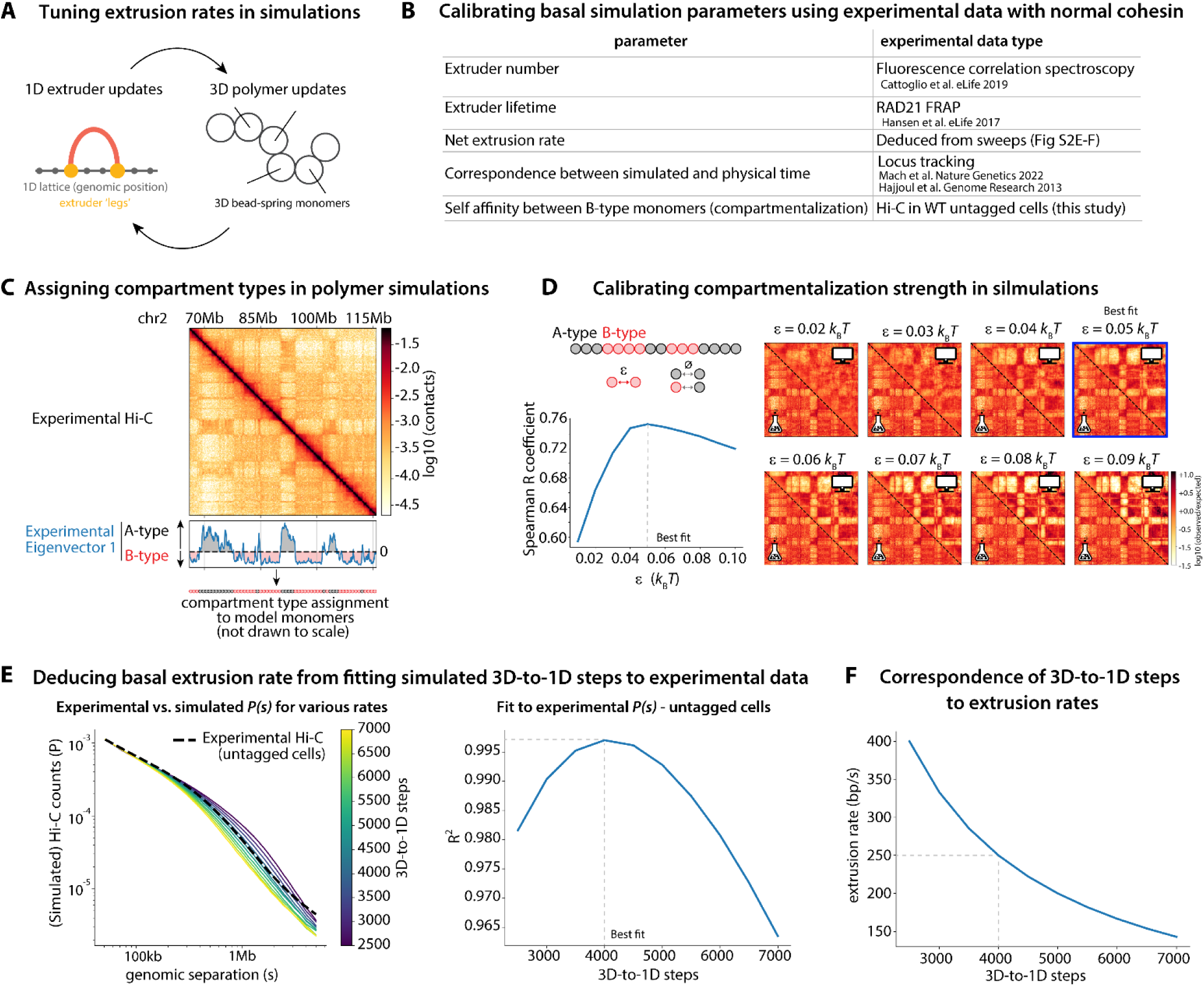
Additional information about modeling approaches. (A) Extrusion simulations alternate between updating positions of polymer beads in 3D and extruder leg positions in 1D (along the chromosome). Tuning extrusion rates in simulations requires modification of the distance traveled and the relative relaxation rate of the polymer fiber. (B) Experimental data used to calibrate simulations. (C) Illustration of how compartment identity is assigned in simulations. (D) The self-affinity (interaction energy) between B-type monomers in simulation was determined by fitting to experimental Hi-C from untagged control cells (see methods). Heatmaps highlight experimental (lower left corner) v.s simulated (top right corner) over a range of self affinities. Note that with this layout for compartments, compartmentalization had minimal impact on simulated *P(s)* for the range of B-B attraction strengths considered. (E) Sweep of polymer model versus lattice model timescales (i.e., 3D-to-1D steps), where each Molecular Dynamics step was estimated to amount to 5 ms of physical time (methods). 4000 3D-to-1D steps best fit the experimental Hi-C *P(s)* of WT untagged cells. (F) Effective average extrusion rate as a function of 3D-to-1D steps, showing that 4000 3D-to-1D steps in WT untagged cells correspond to an extrusion rate of 250 bp/s.

**Fig. S4:**
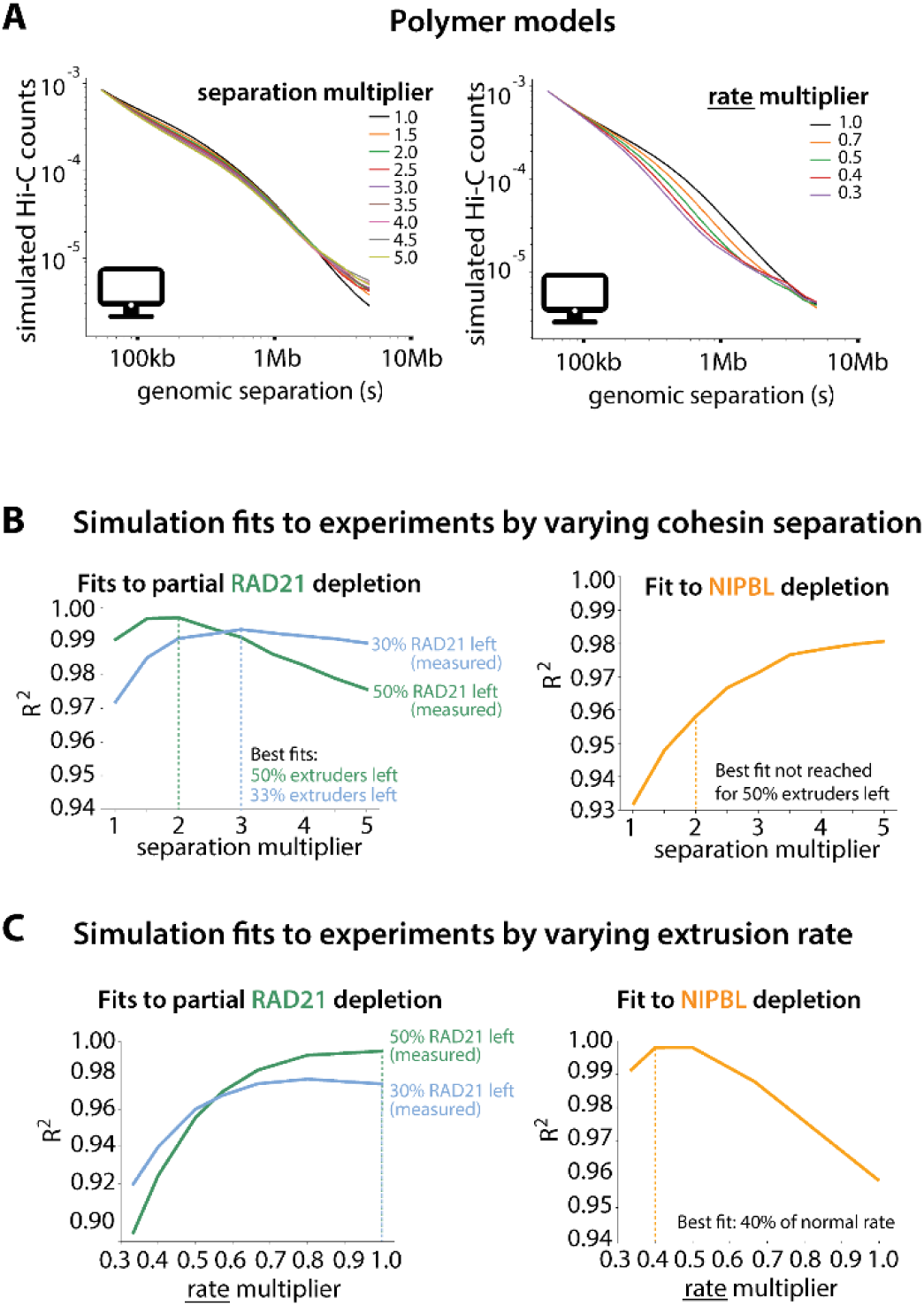
Inferring changes to loop extrusion parameters upon NIPBL depletion or cohesin titration by fitting polymer simulations to experimental Hi-C. (A) in silico exploration of the effect of cohesin separation and extrusion rate on *P(s)* curves from polymer simulations (the basal separation modeling WT untagged cells was set to 185 kb and basal rate was 0.25 kb/sec). (B) Fit scores between simulated *P(s)* curves with varying cohesin separation and experimental Hi-C after partial cohesin (RAD21) depletion (left), or NIPBL depletion (right). For partial cohesin depletion the best models fits closely match the levels of cohesin measured experimentally by flow cytometry. In contrast, for NIPBL-depleted cells the best model fit does not correspond to the experimentally measured amount of bound cohesin. (C) Fit scores as in (B) but varying the extrusion rate. Lowering extrusion rates decreases the fit to partial cohesin depletions. In contrast, for NIPBL-depleted cells lowering extrusion rates increase the fit with experimental Hi-C. Genome folding defects in NIPBL-depleted cells are therefore best explained by slower extrusion rates, rather than lower cohesin loading.

**Fig. S5:**
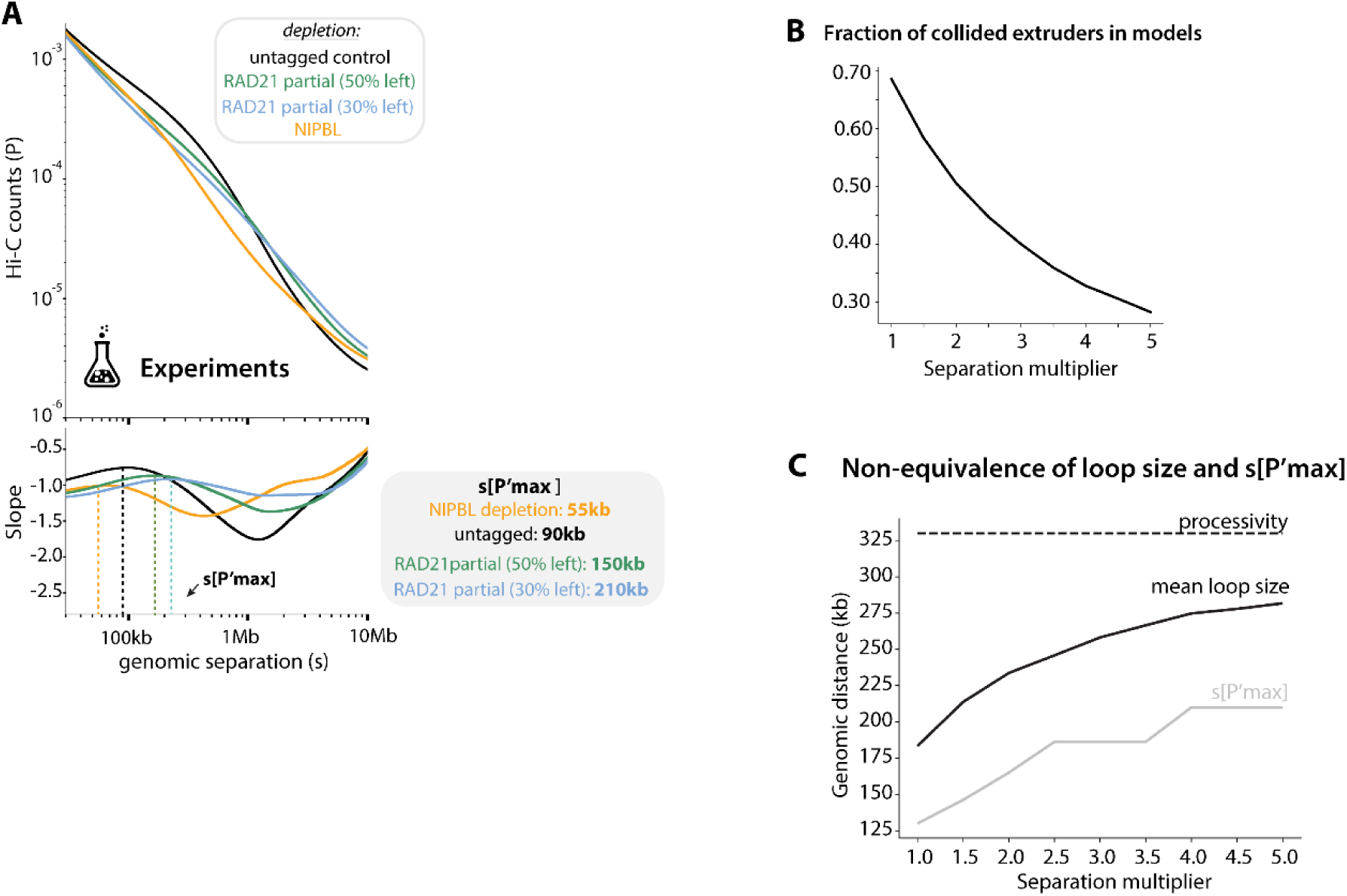
Analysis of *P(s)* derivatives and relationship between s[P’max] and DNA loop size. (A) s[P’max] is defined as the inflection point of the Hi-C *P(s)*, or the first maximum of its slope. (B) Scaling of mean loop size and s[P’max] according to extruder separation in simulations. (C) The simulated fraction of extruders with at least one collided side diminishes as the separation between them increases. Collisions between cohesins therefore limit loop growth, explaining why reducing cohesin levels to 50% or 30% increases loop size.

**Fig. S6:**
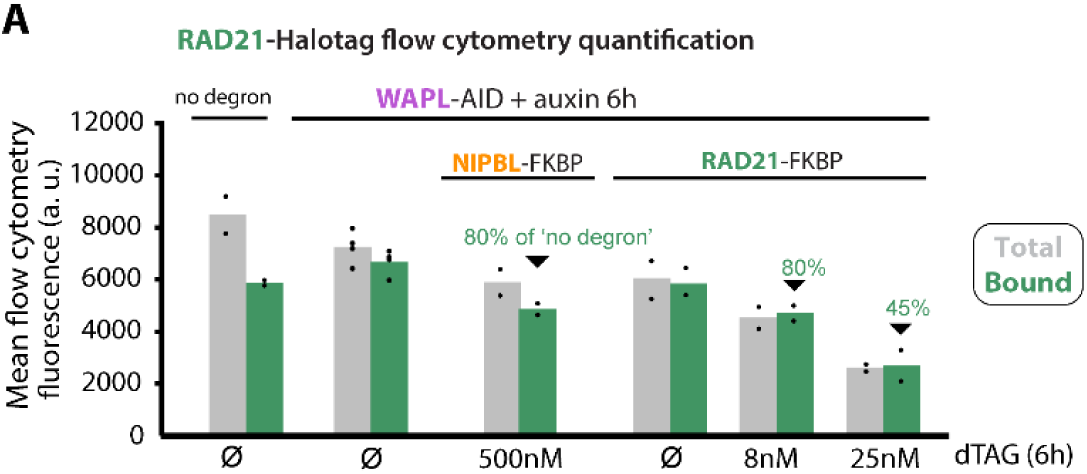
RAD21-Halotag quantifications in WAPL-depleted cells. (A) Flow cytometry quantification of RAD21, highlighting the conditions used for Hi-C.

**Fig. S7:**
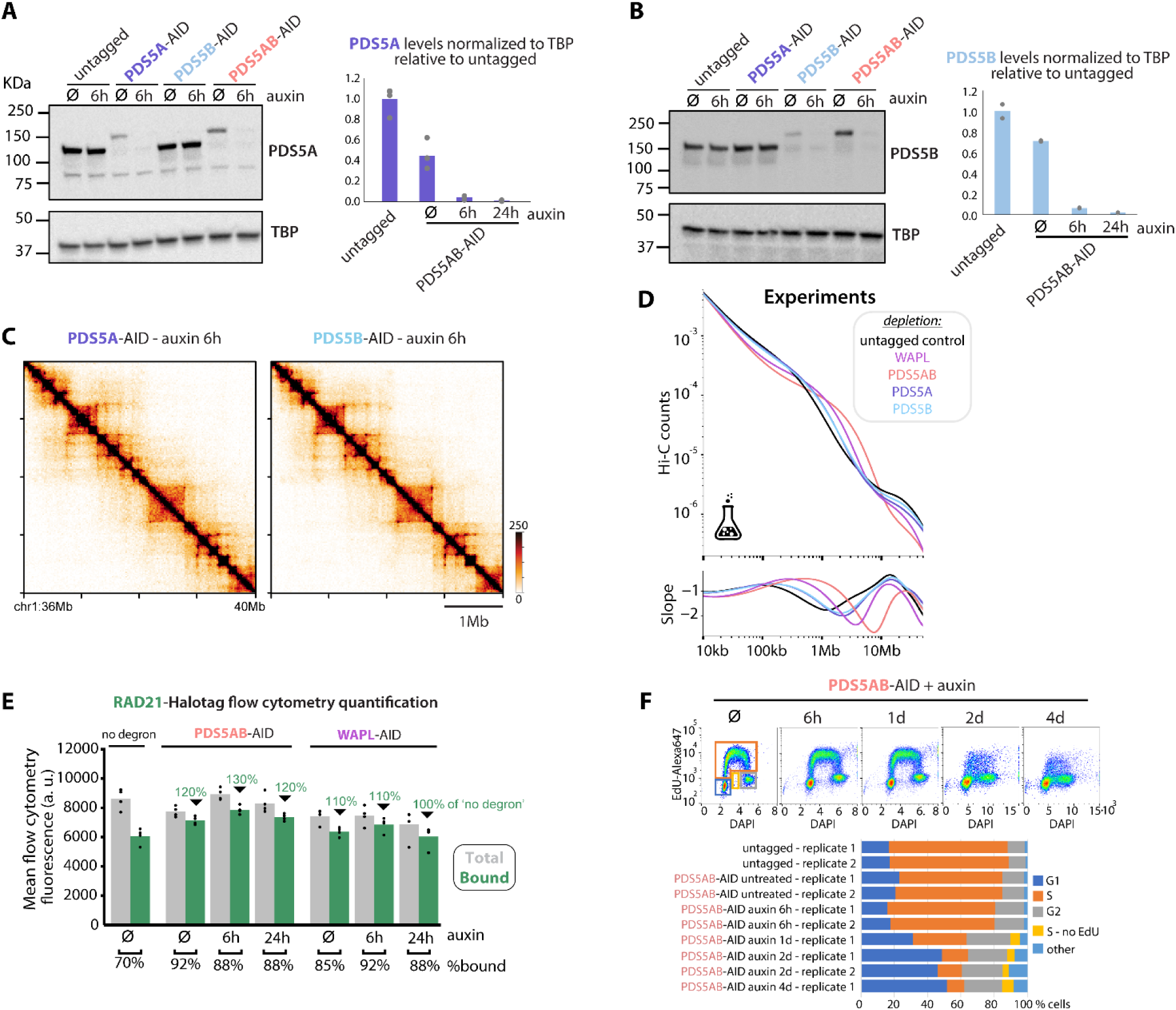
Additional characterization of PDS5 degron cells. (A and B) Western blot quantification of PDS5A and PDS5B depletion. (C and D) Experimental Hi-C (C) and corresponding *P(s)* curves (D) after depleting either PDS5A and PDS5B, highlighting that each paralog contributes to overall chromosome folding in ESCs. (E) Flow cytometry quantification of RAD21-Halotag after PDS5AB or WAPL depletion. Degron leakiness of the AID system, as seen in (A), accounts for the elevated chromatin-bound fraction prior to adding auxin in PDS5AB-AID or WAPL-AID cells. (F) PDS5AB depletion does not cause cell cycle defects within the first day, but longer depletions result in replication stress.

**Fig. S8:**
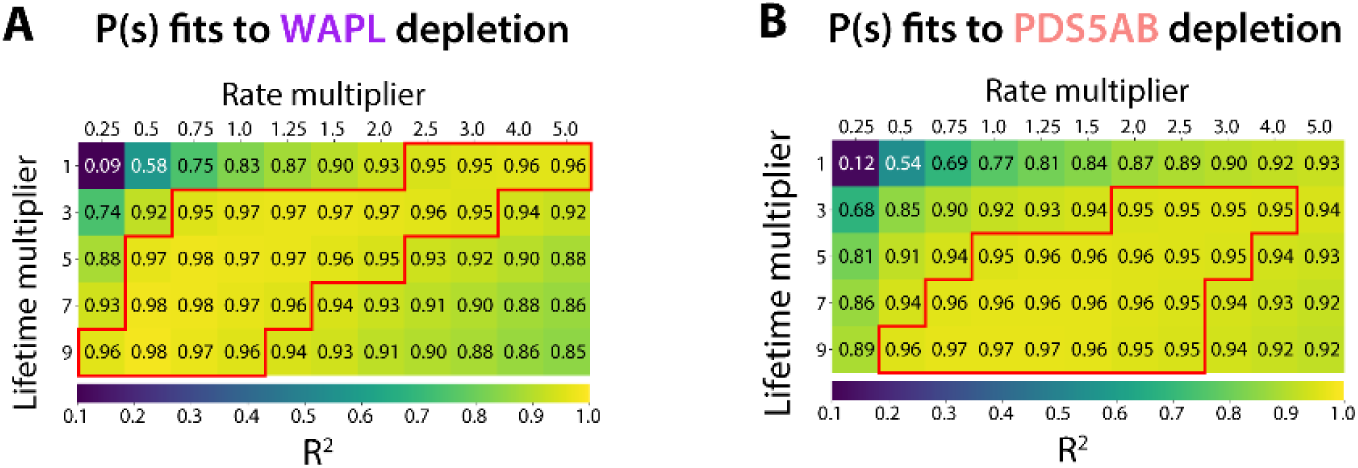
Fits between Hi-C after WAPL or PDS5AB depletion and polymer models across a range of lifetime and rate values. (A) Goodness-of-fit between experimental and simulated *P(s)* as a function of lifetime and extrusion rate after WAPL depletion, or (B) PDS5AB depletion. Red boxes highlight that many combinations of these parameters produce good fits (R^2^>0.95, red boxes) to the experimental *P(s)*. The basal lifetime and rate modeling WT untagged cells was set to 22 minutes and 0.25 kb/sec respectively).

**Fig. S9:**
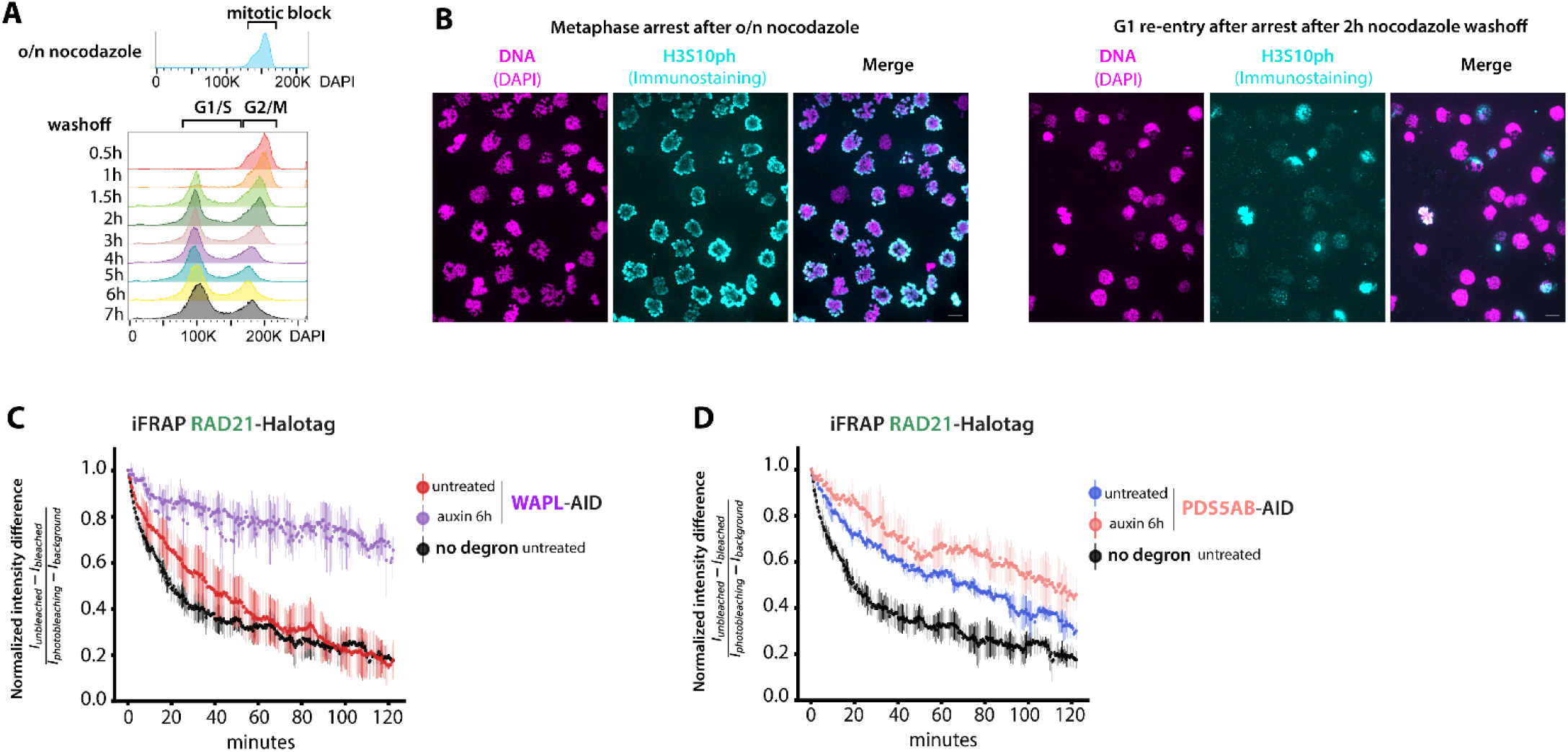
Additional characterizations of WAPL and PDS5AB degron cells. (A) Synchronization controls of DNA content (DAPI) measured by flow cytometry after nocodazole-induced mitotic block. Note that only a fraction of cells re-enter G1 after nocodazole washoff, precluding any bulk assays with this setup. (B) Microscopy examples of unreleased cells blocked in M-phase (left), or after 2h of nocodazole washoff (right). Cells that have not re-entered G1 are eliminated from downstream vermicelli analyses based on their high H3S10Ph immunostaining. (C-D) Inverse fluorescence recovery after photobleaching (iFRAP) of RAD21-Halotag after WAPL of PDS5AB depletion with untreated samples. Lifetime increase in PDS5AB-AID cells prior to auxin treatment can be attributed to leakiness of the degron. n=12-18 cells. Shown are medians +/- standard deviation.

**Fig. S10:**
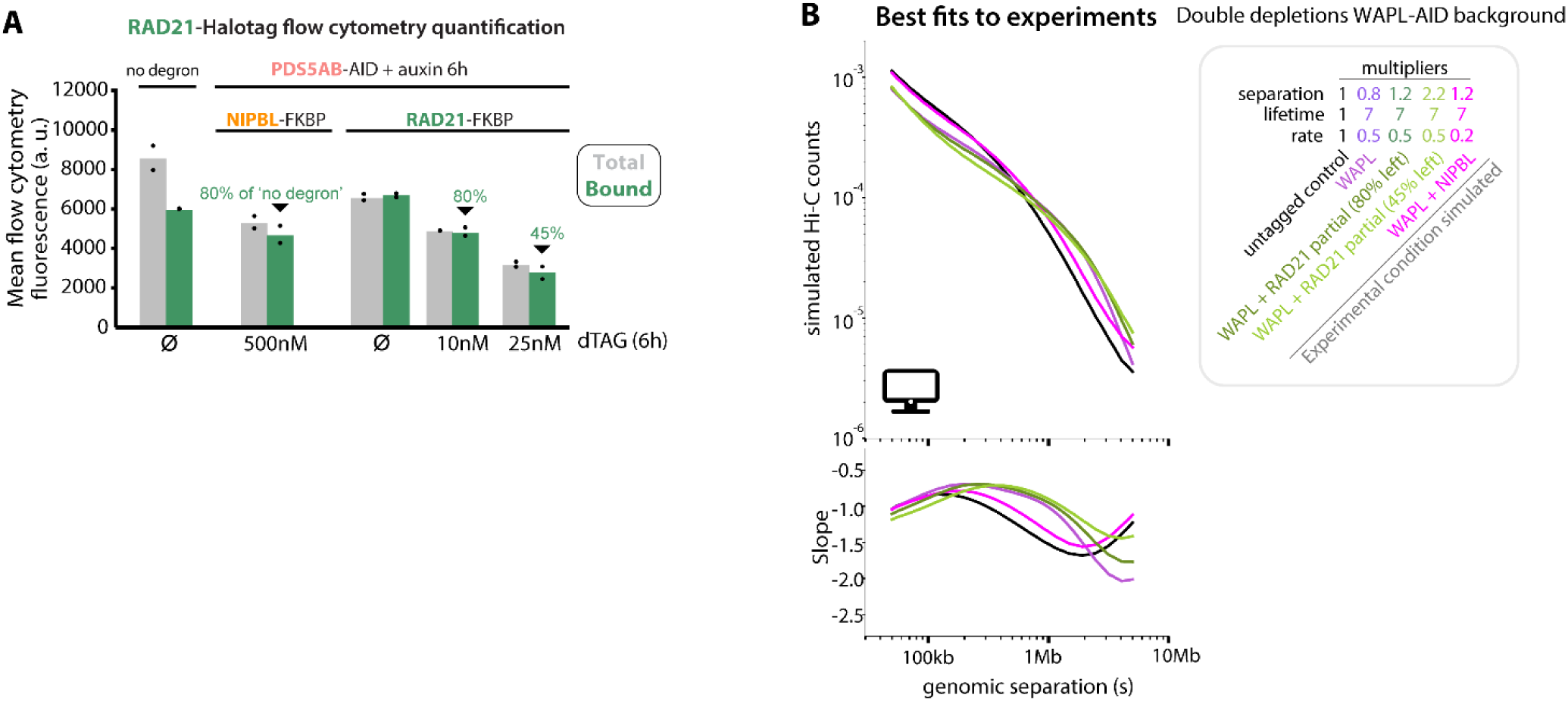
Additional characterization of degron cells. (A) Flow cytometry quantification of RAD21-Halotag, highlighting the conditions used for Hi-C in the indicated PDS5AB-AID backgrounds. (B) Simulated *P(s)* curves and corresponding parameters of loop extrusion as in fig. 5, but in the WAPL-depletion background. These simulations contrast the effect of co-depleting NIPBL or RAD21 together with WAPL, highlighting how an increase in lifetime (WAPL depletion) can be offset by a decrease in rate (NIPBL depletion) - yielding *P(s)* that resemble the basal situation in untagged cells. Lifetime increase cannot be offset by reducing the amount of extruders (partial RAD21 depletion).

**Fig. S11:**
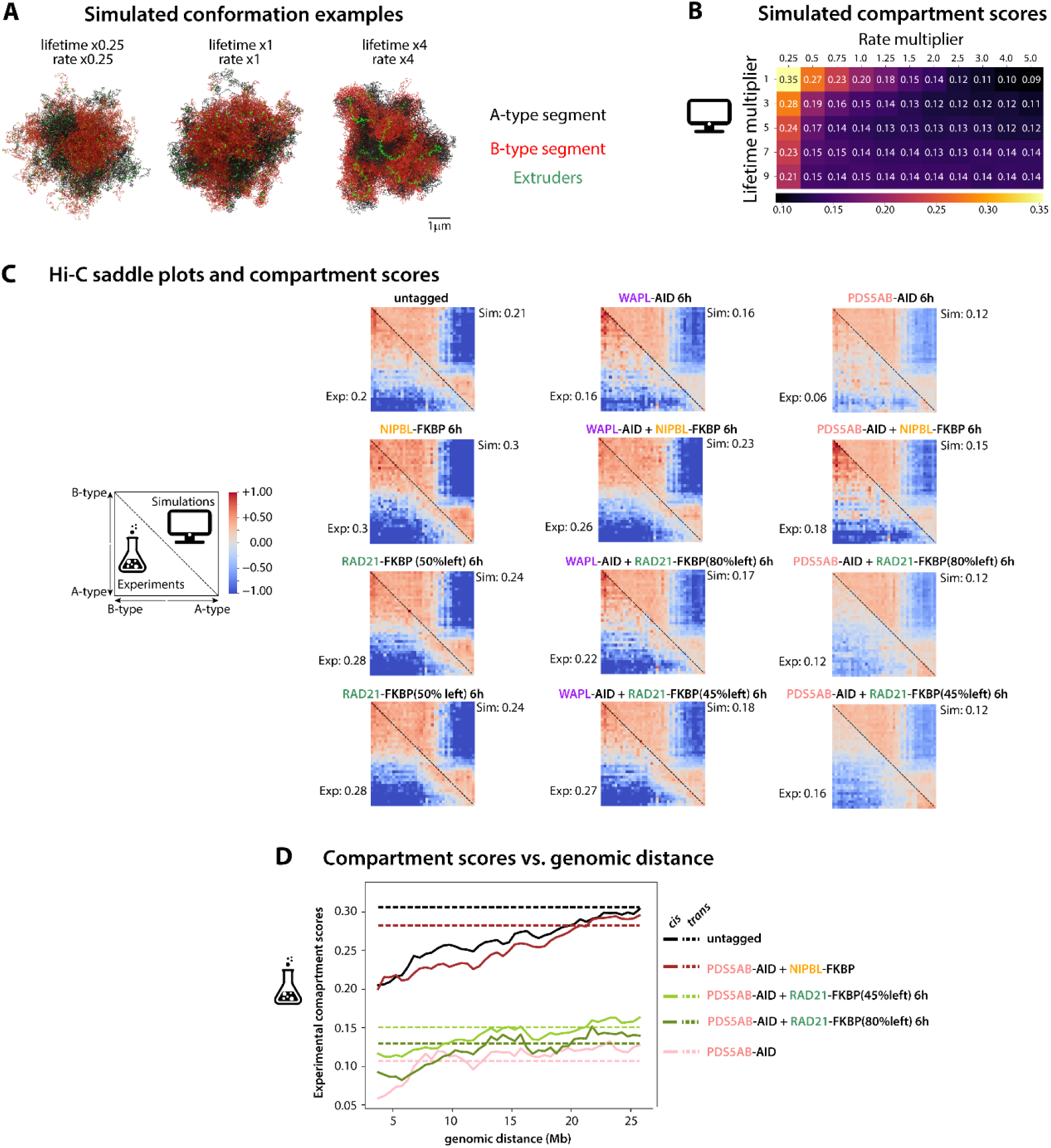
Experimental and simulated compartmentalization defects across loop extrusion mutants and regimes. (A) Example of simulated conformations highlighting the spatial positions of compartment types and extruders. (B) Simulated compartment scores obtained with a range of lifetime and rate multipliers, at basal extruder separation. (C) Comparison of Hi-C saddle plots obtained from simulations and experiments with corresponding compartment scores. (D) Experimental compartment scores from Hi-C across genomic distances in *cis* and in *trans* (see methods).

**Fig. S12:**
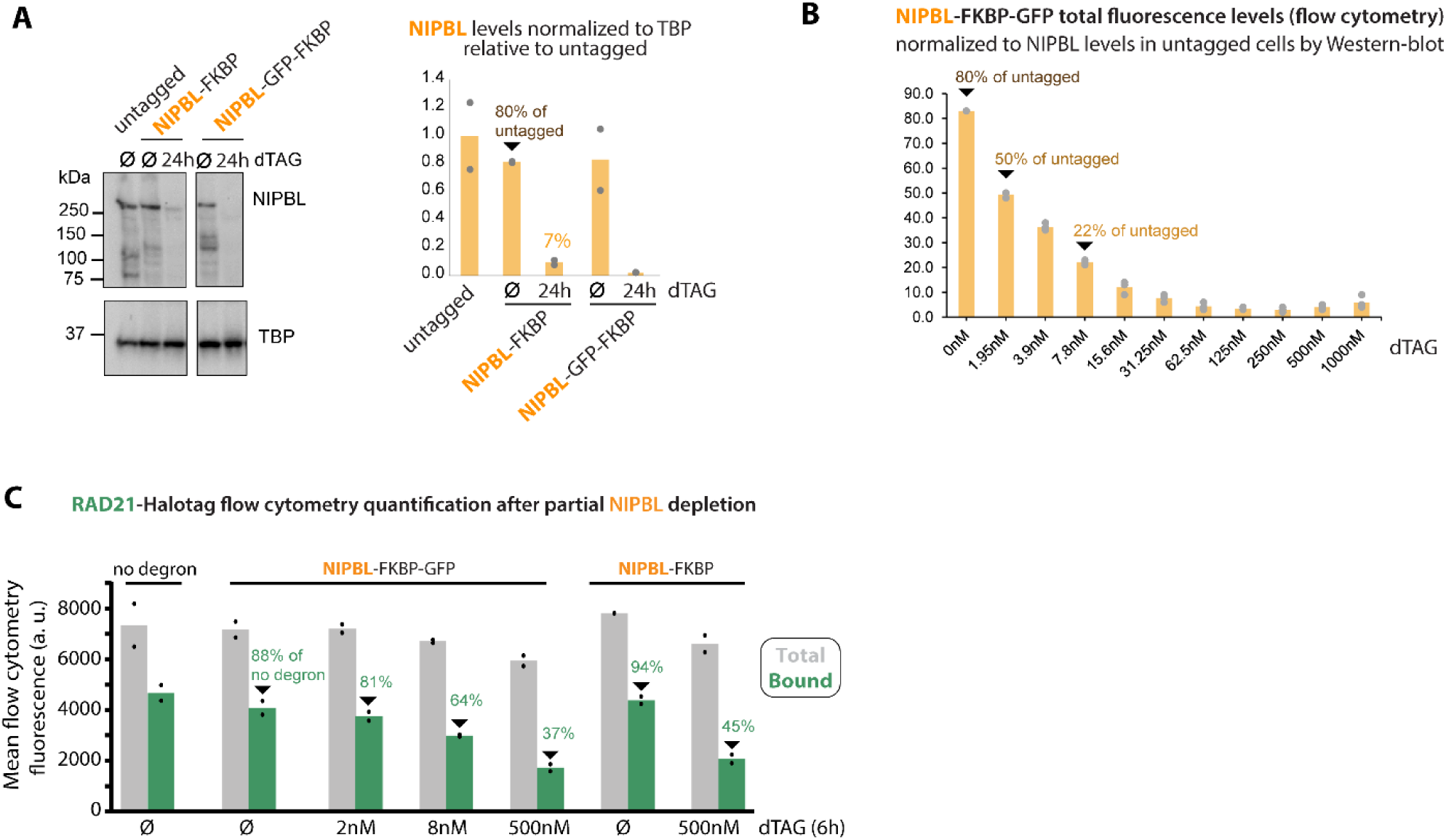
Quantification of cohesin and NIPBL in cells titrated for NIPBL. (A) Western blot quantification of basal NIPBL levels in NIPBL-FKBP and NIPBL-FKBP-GFP ESCs, and confirmation of degron potency with 500mM dTAG (maximum depletion). (B) NIPBL-FKBP-GFP levels after dTAG titration, normalized to basal levels in untagged cells as estimated by Western-blots. Doses used for Hi-C are indicated. (C) Flow cytometry quantification of RAD21-Halotag, highlighting the conditions used for Hi-C and corresponding simulations in the indicated NIBPL-degron cells.

**Video S1:**
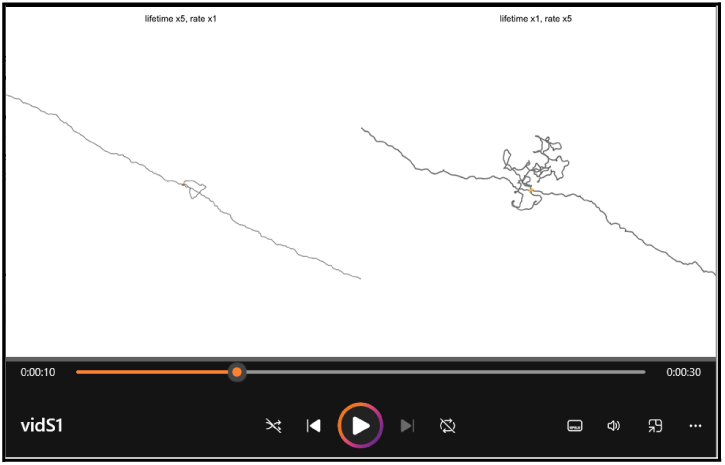
vidS1.mp4 - Differential effects of chromatin lifetime and extrusion rate on single-cohesin behavior. Representative simulation time-lapse (sped up x50 relative to physical time) of extrusion by isolated cohesin at baseline rate and increased lifetime (left), or baseline lifetime and increased rate (right). The chromatin fiber is artificially stretched by an external force of magnitude ∼0.5 pN applied at the chain extremities to facilitate visualization.

**Video S2:**
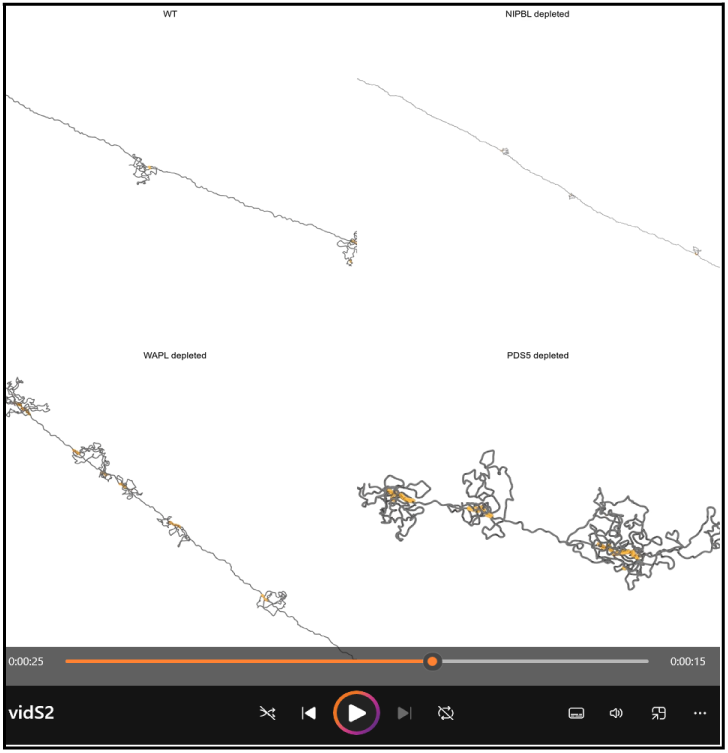
vidS2.mp4 - Simulated impacts of extrusion parameters on local chromatin structure and dynamics. Same as Video S1 for a stretched 2.5 Mb chromatin segment subjected to extrusion at either baseline (top left) or altered cohesin separations, lifetimes and rates, as respectively inferred for NIPBL- (top right), WAPL- (bottom left) and PDS5-depleted mutants (bottom right). See fig. 5G for the full list of parameter values used.

**Table 1: Cell-Lines-Vectors-sgRNAs_v1.xlsx**

**Table 2: Supp_Table_2_HiC_mapping_statistics.xlsx**

**Archive 1: Annotated plasmid maps.zip**

